# USP1/UAF1 targets polyubiquitinated PCNA with an exo-cleavage mechanism that enriches for monoubiquitinated PCNA

**DOI:** 10.1101/2024.12.16.628697

**Authors:** Niels Keijzer, Jan Sakoltchik, Kaustav Majumder, Nina van Lil, Farid El Oualid, Alexander Fish, Titia K. Sixma

**Affiliations:** Netherlands Cancer Institute and Oncode Institute, Plesmanlaan 121, 1066 CX Amsterdam, The Netherlands; UbiQ Bio B.V., Science Park 301, 1098 XH, Amsterdam, The Netherlands

## Abstract

DNA damage tolerance (DDT) is an important pathway that allows our cells to bypass DNA lesions during replication. DDT is orchestrated by ubiquitination of PCNA: Monoubiquitination (PCNA-Ub) initiates recruitment of TLS polymerases but also serves as substrate for K63-linked polyubiquitination that leads to HR-mediated bypass mechanisms. Recent work on USP1/UAF1 inhibition revealed that K48-linked chains are also formed on PCNA, resulting in its proteasomal degradation. USP1/UAF1 is established as deubiquitinating enzyme (DUB) for PCNA-Ub, but little is known about deubiquitination of chains on PCNA.

Here we show that USP1/UAF1 cleaves both K48 and K63-linked ubiquitin chains on PCNA efficiently, using an exo-cleavage mechanism. Kinetic analysis reveals that USP1/UAF1 prefers cleaving the ubiquitin-ubiquitin bond over cleavage of the ubiquitin-PCNA isopeptide bond and therefore treats poly- and monoubiquitinated PCNA as different substrates. A cryo-EM structure of USP1/UAF1 with a K63-diubiquitin and structure-based mutagenesis reveals that its mechanistic preference is maintained in evolution. Its kinetic mechanism results in relative enrichment of monoubiquitinated PCNA that could initially promote TLS over HR-like bypass. Taken together, these results suggest that USP1/UAF1 could be important in temporary protection of PCNA against K48- and K63-linked polyubiquitination and highlight this DUB as a potential regulator of DDT pathway choice.

## Introduction

Ubiquitination plays a crucial role in regulation of the response to DNA damage (Al-Hakim et al., 2010). There is extensive variety in ubiquitin signalling: Ubiquitin can be attached as a single entity or form different types of polyubiquitin chains through linkage to any of ubiquitin’s seven internal lysines or N-terminal methionine (Hershko & Ciechanover, 1998; Komander & Rape, 2012). Each of these different types of ubiquitination can lead to different outcomes on a protein substrate. Reversion of ubiquitination by deubiquitinating enzymes (DUBs) is therefore a crucial component in ubiquitin signalling (Komander et al., 2009; Nijman, Luna-Vargas, et al., 2005).

USP1 is a DUB with important roles in the response to DNA damage and is emerging as a promising clinical target (Cadzow et al., 2020; Simoneau et al., 2023). USP1 functions as a heterodimer with its essential cofactor UAF1, which hyperactivates USP1 by relieving autoinhibition caused by USP1’s catalytic domain insertions 1 and 3 (Dharadhar et al., 2021). USP1/UAF1 is a highly specific DUB, which targets PCNA and FANCI/FANCD2 (Huang et al., 2006; Nijman, Huang, et al., 2005). This gives USP1/UAF1 important roles in regulation of DNA damage tolerance (DDT) through action on PCNA and in interstrand crosslink (ICL) repair through action on Fanconi anaemia proteins FANCI/FANCD2.

During replication, PCNA and its interacting polymerase encounter numerous DNA lesions, which block the replicative polymerase from inserting the next base. In turn, this can lead to replication fork stalling, fork collapse and overall genomic instability. To prevent this, DDT enables bypass of DNA lesions instead of waiting for DNA repair mechanisms, which allows for swift continuation of replication and maintenance of genome stability. Alternatively, repriming mediated by PrimPol can ensure that replication continues downstream of the lesion (Bainbridge et al., 2021), but this leaves a ssDNA gap behind the stalled PCNA, which subsequently requires gap filling by DDT (Nusawardhana et al., 2024; Piberger et al., 2020; Tirman et al., 2021).

DDT is initiated by stalling of PCNA at the DNA lesion, which sets off a cascade of reactions that culminates in monoubiquitination of PCNA at its K164-residue (PCNA-Ub) by Rad6/Rad18 (Hoege et al., 2002). PCNA-Ub exchanges its bound replication polymerase for a translesion synthesis (TLS) polymerase (Bienko et al., 2005; Kannouche et al., 2004; Lehmann et al., 2007). These polymerases can bypass DNA lesions due to their lack of proofreading activity and modified active sites that can accommodate different types of lesions (Biertümpfel et al., 2010; Kirouac & Ling, 2011). While these features render the TLS polymerases more error-prone, TLS can be coupled with mismatch repair to correct for wrongly inserted bases (Tsaalbi-Shtylik et al., 2024). PCNA-Ub is also a substrate for K63-linked polyubiquitination (Hoege et al., 2002), which is catalyzed by a different set of E2 and E3 enzymes. These have been identified as Ubc13/Mms2 and Rad5 in yeast (Hoege et al., 2002; Parker & Ulrich, 2009), while in humans multiple E3s have been identified (Gallina et al., 2021; Krijger et al., 2011; Unk et al., 2006, 2008).

Polyubiquitinated PCNA (K63-PCNA-Ub_N_) leads to two homologous recombination (HR)-mediated pathways, referred to as template switching (TS) and fork reversal (FR), that can also bypass DNA lesions (Hoege et al., 2002; Stelter & Ulrich, 2003). Both pathways use the undamaged sister strand as a template and are considered to be error-free (Chiu et al., 2006). However, what exactly triggers formation of K63-PCNA-Ub_N_ and why HR-mediated bypass is required along with TLS is not clear. Additionally, it is unclear when fork reversal and template switching are preferred over translesion synthesis. It is possible that these HR-mediated bypass mechanisms are required for bulkier lesions which would not fit in the TLS polymerase active site (Sale et al., 2012).

Timely removal of ubiquitin is important since it allows for continuation of regular replication. Deubiquitination of PCNA-Ub is carried out by USP1/UAF1 (Huang et al., 2006) and has been studied extensively. USP1/UAF1 can inhibit itself by autocleavage which happens upon exposure to UV-light (Huang et al., 2006), which promotes DDT. It is thought that TLS polymerases are required to extend about 5-60bp after the lesion (Fujii & Fuchs, 2004), after which a different USP1/UAF1 comes to deubiquitinate PCNA, allowing a replicative polymerase to take over again. Deubiquitination of K63-PCNA-Ub_N_ is less well established and it is still unclear what triggers formation of these chains on PCNA. While USP1/UAF1 activity against K63-linked chains on PCNA was reported once (Yang & Zou, 2009), such a role has never been studied in detail.

Recent studies on USP1 inhibition demonstrated the accumulation of polyubiquitinated PCNA upon loss of USP1 (Cadzow et al., 2020; Simoneau et al., 2023). Further analysis showed that in absence of functional USP1, K48-linked chains accumulate on PCNA which leads to degradation by the proteasome. This K48 chain formation on PCNA is accomplished by UBE2K and RNF138 and requires monoubiquitination of PCNA by RAD6/RAD18 (Simoneau et al., 2023). Inhibition of USP1 is especially effective in BRCA1/2 deficient tumours, as it is synthetically lethal with PARP inhibition in the absence of BRCA1/2, and helps to overcome PARP resistance (Cadzow et al., 2024; Lim et al., 2018; Simoneau et al., 2023). This shows that replication stress caused by HR-deficiency requires PCNA-Ub or K63-PCNA-Ub_N_ mediated bypass. It is unclear whether other types of replication stress also trigger K48-linked ubiquitination of PCNA and under what circumstances K48-mediated degradation of PCNA occurs.

DUBs process ubiquitin chains using different cleavage patterns. They can cleave the entire chain off the substrate (base cleavage), cleave a ubiquitin-ubiquitin linkage in the middle of the chain (endo-cleavage), or start at the distal ubiquitin and processively cleave the chain one ubiquitin at a time (exo-cleavage) (Fig. 1A). In context of PCNA ubiquitination, base cleavage immediately creates free PCNA, while endo- or exo-cleavage can leave either monoubiquitinated or polyubiquitinated PCNA, which in turn affects the DDT outcome.

**Figure 1.**
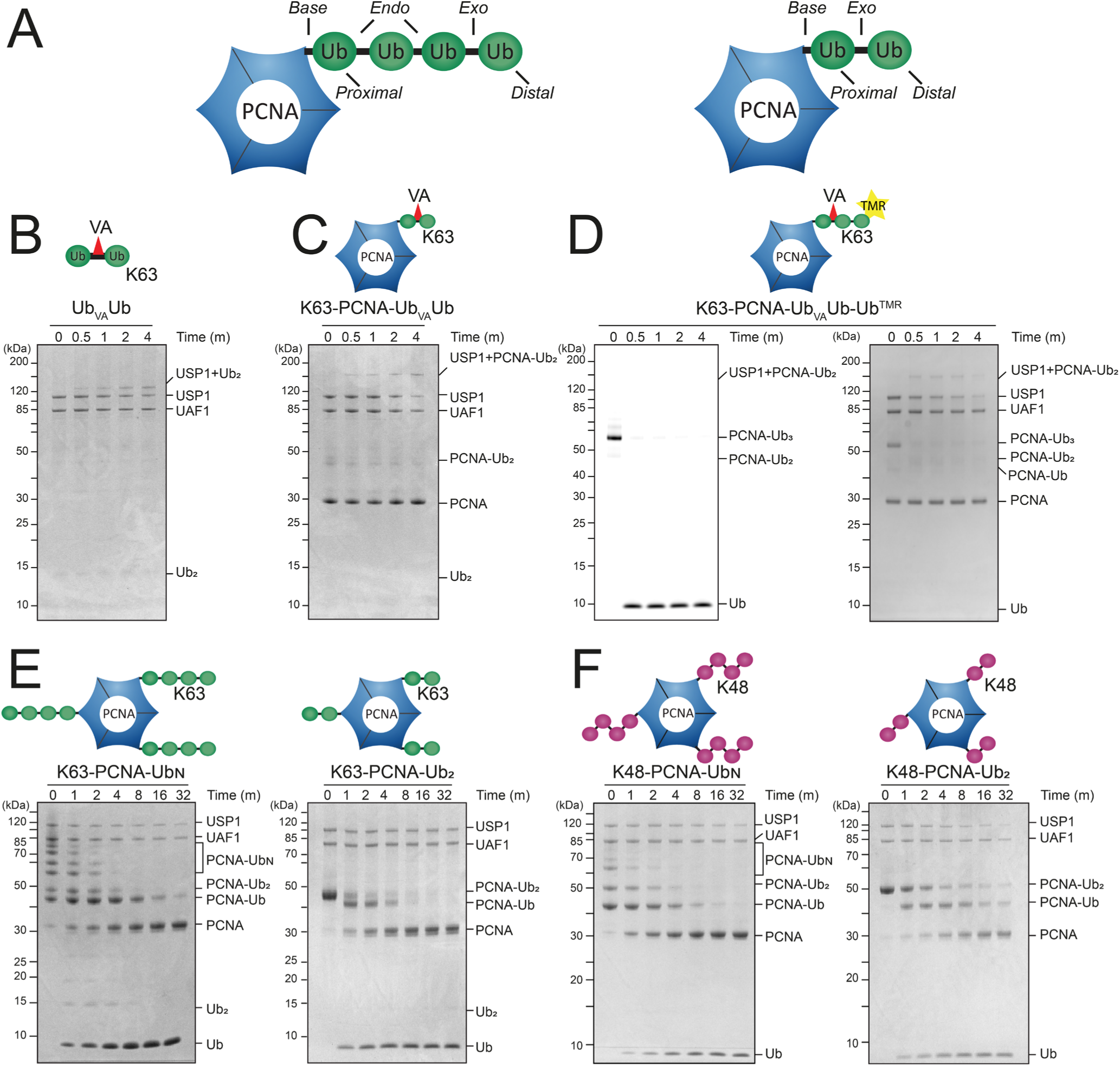
USP1/UAF1 strongly prefers exo-cleavage on K63- and K48-polyubiquitinated PCNA. A) Schematic of cleavage patterns on PCNA-Ub_N_ and PCNA-Ub_2_ B-E) SDS PAGE gels from DUB activity assays B) Conjugation reaction of USP1/UAF1 to Ub_VA_Ub in a complex of USP1 with a Ub_2_ chain. C) Conjugation of USP1/UAF1 to K63-PCNA-Ub_VA_Ub shows a large band shift corresponding to USP1 performing exo-cleavage. D) Fluorescence scan (left panel) and Coomassie-stained SDS PAGE gel (right panel) of K63-PCNA-Ub_VA_Ub-Ub_TMR_ and USP1/UAF1 reaction shows fast release of Ub^TMR^, and subsequent USP1 conjugation to PCNA-Ub_VA_Ub only visible on the Coomassie stained gel. E) DUB assay of USP1/UAF1 on K63-PCNA-Ub_N_ (left panel) and K63-PCNA-Ub_2_ (right panel) shows rapid disappearance of chains, enrichment of PCNA-Ub and appearance of a free mono-ubiquitin, indicating that USP1/UAF1 prefers exo-cleavage on both longer K63-chains and diubiquitinated PCNA. F) DUB assay of USP1 on K48-PCNA-Ub_N_ (left panel) and K48-PCNA-Ub_2_, shows that USP1 has an even stronger preference for exo-cleavage on K48-chains.

Here we analyze the USP1/UAF1 cleavage mechanism on K63- and K48-linked polyubiquitinated PCNA to understand the cleavage preference and kinetics of this reaction. We find that USP1/UAF1 has a strong preference for exo-cleavage on polyubiquitinated PCNA. Kinetic modeling reveals that USP1/UAF1 is efficient at cleaving both K48- and K63-linked polyubiquitin chains on PCNA and prefers cleaving Ub-Ub bonds over the Ub-PCNA bond. By solving the cryo-EM structure of active full-length USP1/UAF1 covalently bound to a K63-linked diubiquitin chain, we show how USP1/UAF1 binds to K63-ubiquitin chains. Structure-function analysis reveals that UAF1 plays a minor role USP1/UAF1’s exo-cleavage preference and that the preference for Ub-Ub is caused primarily by USP1. The result of this preference is relative enrichment of monoubiquitinated PCNA. This could be important, as it would promote TLS over HR-like mediated bypass. These results show how USP1/UAF1 can act as an important regulator of DDT and how it can direct DDT pathway choice.

## Results

### USP1/UAF1 prefers to cleave K63- and K48-PCNA-UbN using exo-cleavage

To study the cleavage mechanism of USP1/UAF1, we produced a variety of defined K63- or K48-ubiquitinated PCNA substrates (Fig. S1), using wildtype ubiquitin, fluorescently labeled ubiquitin and activity based ubiquitin probes. Together, these substrates help us to precisely follow deubiquitination of polyubiquitinated PCNA by USP1/UAF1. We used PCNA with defined monoubiquitination on K164, using established enzymatic modification (Hibbert & Sixma, 2012), and extended these to diubiquitin and/or longer ubiquitin chains with either K63 or K48 specific E2 enzymes and a chain-promoting E3 ligase, RNF8. For kinetic analysis we used TAMRA (TMR)-labeled ubiquitin, covalently attached to an A28C-mutated ubiquitin. This allowed PCNA to be successfully modified with TMR-labeled Ub^A28C^ by the E1, E2s and E3 used in this study and did not interfere with USP1/UAF1 cleavage (Fig. S2).

Additionally, we used a diubiquitin with a VME-like warhead mimicking the K63 linkage (K63-linked diubiquitin vinyl amide (VA), here termed Ub_VA_Ub) (Mulder et al., 2014), which conjugates to USP1 after USP1/UAF1 attempts to cleave the Ub-Ub bond (Fig. 1B, Fig. S1). To examine the USP1/UAF1 cleavage mechanism on PCNA (Fig. 1A), we then enzymatically attached this diubiquitin probe to PCNA’s K164 residue, aiming for one diubiquitin probe per PCNA trimer. This allows us to distinguish between cleavage of the PCNA-Ub bond (base-cleavage) or the Ub-Ub bond (exo-cleavage) by following the band shift after USP1 attempts to cleave K63-PCNA-Ub_VA_Ub. Addition of USP1/UAF1 results in conjugation to K63-PCNA-Ub_VA_Ub (Fig. 1C), revealing that USP1 performs exo-cleavage.

We then generated triubiquitinated PCNA by attaching a TAMRA labeled Ub^K63R^ to K63-PCNA-Ub_VA_Ub. This yields K63-PCNA-Ub_VA_Ub-Ub^TMR^, a substrate that allows distinction between exo-, endo- and base-cleavage by following Ub^TMR^. On this substrate, cleavage by USP1/UAF1 results in formation of free Ub^TMR^, while K63-PCNA-Ub_VA_Ub-Ub^TMR^ is almost completely consumed within one minute (Fig. 1D), which can only happen if USP1/UAF1 uses an exo-cleavage mechanism. On the Coomassie stained gel, we observe conjugation of USP1 to K63-PCNA-Ub_VA_Ub, demonstrated by the same 40 kDa shift of USP1 as seen in Figure 1C, in the next step of exo-cleavage. USP1 does not conjugate to K63-PCNA-Ub_VA_Ub-Ub_TMR_, indicating that USP1 exclusively uses exo-cleavage in the first turnover and thus has a strong preference for trimming ubiquitin chains on PCNA.

We then performed deubiquitination assays on native K63-linked polyubiquitinated PCNA (K63-PCNA-Ub_N_) and K63-linked diubiquitinated PCNA (K63-PCNA-Ub_2_). On K63-PCNA-Ub_2_, USP1/UAF1 cleaves the ubiquitin-ubiquitin isopeptide-bond (exo-cleavage), as we can observe that PCNA-Ub and free monoubiquitin are being produced (Fig. 1E, right panel). On longer ubiquitin chains (K63-PCNA-Ub_N_), USP1/UAF1 appears to favor exo-cleavage over endo- or base-cleavage, as it primarily produces free monoubiquitin (Fig. 1E, left panel). The appearance of minor amounts of free ubiquitin chains of different lengths indicates that some base- or endo-cleavage takes place as well. Endo-cleavage is most likely, since USP1/UAF1 does not use base-cleavage on PCNA-Ub_2_ either (Fig. 1E, right panel). These free ubiquitin chains are further cleaved by USP1/UAF1 and are completely consumed after 8 minutes, which is also when most K63-PCNA-Ub_N_ has been processed. Interestingly, USP1/UAF1 appears to prefer cleaving K63-PCNA-Ub_N_ over monoubiquitinated PCNA, as demonstrated by PCNA-Ub bands increasing until t = 4 minutes and decreasing only when polyubiquitinated PCNA has depleted.

Cleavage of K48-linked chains on PCNA by USP1/UAF1 (Fig. 1F) is equivalent to K63 chains and shows that USP1/UAF1 exclusively employs exo-cleavage on this chain type. Chains shorten over time and only free ubiquitin is produced. As was observed for K63-PCNA-Ub_N_ USP1/UAF1 starts cleaving monoubiquitinated PCNA only after the majority of chains has been processed, suggesting that USP1/UAF1 prefers cleaving Ub-Ub bonds over the Ub-PCNA bond in K48-PCNA-Ub_N_ as well.

### Global kinetic analysis

In order to further characterize the cleavage reaction employed by USP1/UAF1, we set out to generate a global kinetic model. We used K63- and K48-PCNA-Ub_2_, where either the proximal or distal ubiquitin was fluorescently labeled with TAMRA (TMR), along with PCNA-Ub^TMR^. These substrates allowed us to follow USP1/UAF1 activity in fluorescence polarization assays (Fig. 2A, Fig. S3A,B). In addition, we studied binding affinities of USP1/UAF1 for PCNA-Ub, PCNA and free ubiquitin by MST analysis and determined the activity of USP1/UAF1 on a minimal substrate (Ub-Rho) (Fig. S3C,D). All activity assays and the binding data were fitted simultaneously to a single global kinetic model using KinTek explorer (Fig. 2B, Table S1, Fig. S4A,B).

**Figure 2.**
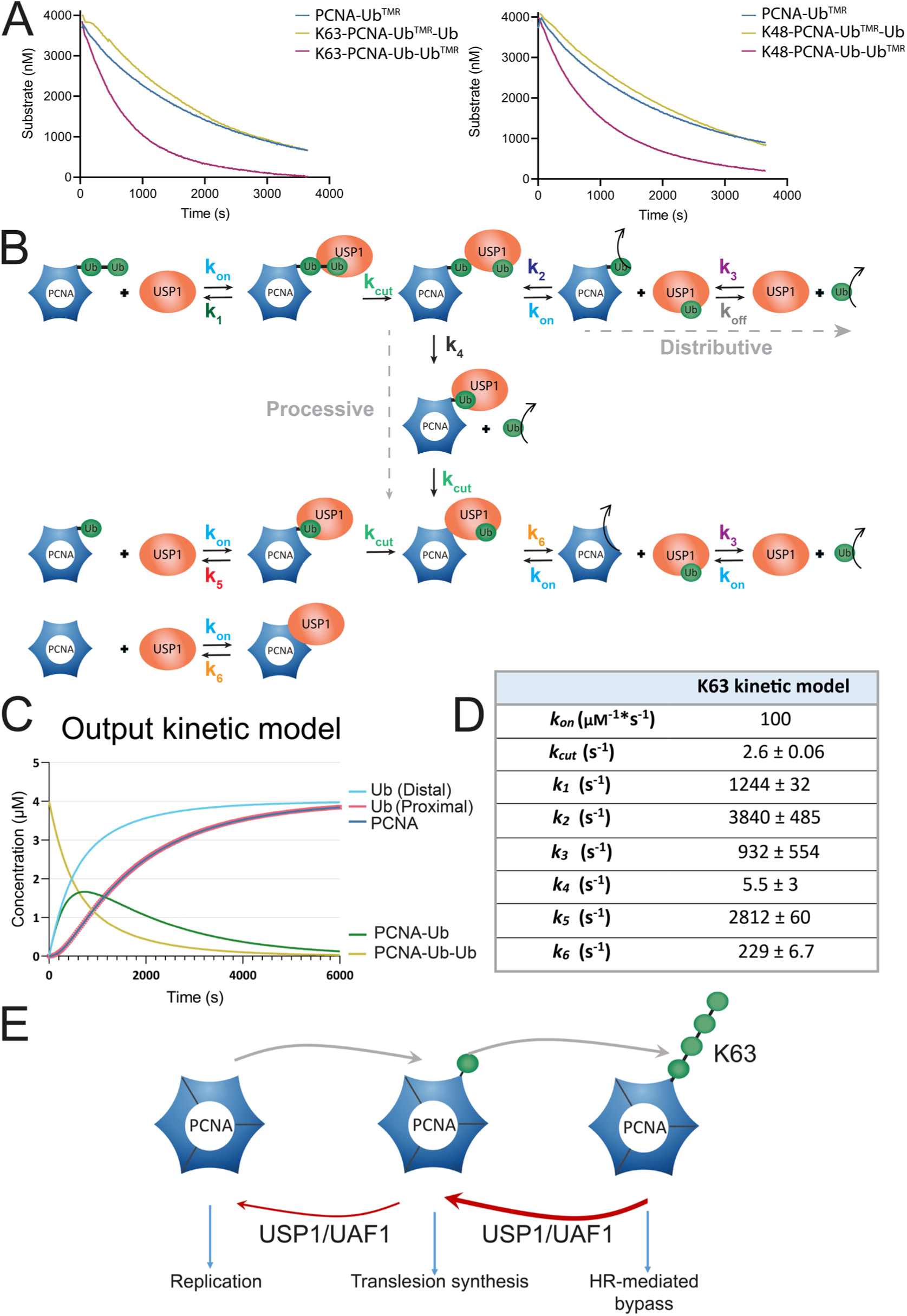
Kinetic modeling of USP1 activity on K48- and K63-polyubiquitinated PCNA. Fluorescence polarization assays of USP1/UAF1 activity comparing a single substrate concentration (4 µM) of PCNA-Ub^TMR^, K63-PCNA-Ub^TMR^-Ub and K63-PCNA-Ub-Ub^TMR^ (left panel) and PCNA-Ub^TMR^, K48-PCNA-Ub^TMR^-Ub and K48-PCNA-Ub-Ub^TMR^ (right panel). The assays reveal that USP1/UAF1 prefers cleaving the ubiquitin-ubiquitin bond of both chain types. Full concentration range is shown in *Supplementary figure 2A*. **B)** Schematic representation of the final Kintek model, which describes deubiquitination of both K63 and K48 substrates. The model shows the potential paths taken by a single entity of USP1/UAF1 (shown as USP1). Written version of the kinetic model and accompanying calculated constants are shown in *Supplementary figure 3*. Since the off rate for PCNA-Ub_2_ (*k_1_*) is threefold lower than the off rate for PCNA-Ub (*K_5_*), USP1 prefers to proceed to another PCNA-Ub_2_ after cleaving and releasing the distal ubiquitin from PCNA-Ub_2_. USP1/UAF1 uses a distributive cleavage mechanism (bind, cleave, release) instead of a processive mechanism, as *k_2_* is much higher than *k_4_.* **C)** Simulation generated by the kinetic model shows cleavage of substrates and appearance of different products as USP1/UAF1 cleaves PCNA-Ub_2_. Its preferred cleavage mechanism and preference for PCNA-Ub_2_ lead to accumulation of PCNA-Ub. **D)** Constants describing the different reactions and equilibriums in the kinetic models. *K_on_* is diffusion limited and therefore the same for each reaction. *k_1_, k_2_*, *k_3_, k_4_, k_5_, k_6_* are the off rates for each reaction and can be seen as representations of affinity in this model. Table including constants of the K48 kintek is shown in Table S1. **E)** USP1/UAF1’s preference for exo-cleavage and cleaving Ub-Ub bonds allows for temporal enrichment of PCNA-Ub, which would promote translesion synthesis over HR-mediated bypass.

To formulate an appropriate model (Fig. 2B, Table S1), we started by keeping options open for exo-, endo- and base-cleavage. Only a model in which USP1/UAF1 was instructed to solely use exo-cleavage resulted in a good fit, which agrees with the exo-cleavage preference observed in our gel based DUB assays (Fig. 1).

An additional important parameter was whether the reaction was processive (bind, cleave and move to the next Ub on the chain), distributive (bind, cleave and release) or both. A model where USP1/UAF1 solely uses processive cleavage gave poor fit to the experimental data. We then wrote a model where USP1/UAF1 solely performs a distributive mechanism, with a bind, cleave and release for each cleavage event, which improved the fit. However, we obtained the best fit when USP1/UAF1 has the option to perform both distributive and processive cleavage. This was interesting because the constants calculated by the final model show that despite using the two mechanisms, USP1/UAF1 has a strong preference for the distributive cleavage mechanism (Table S1).

Important for this distributive cleavage model was that after cleavage of PCNA-Ub_2_, USP1/UAF1 immediately releases PCNA-Ub and remains bound to the distal ubiquitin, rather than PCNA-Ub. Additionally, for this bind, cleave and release mechanism a proper fit could only be obtained if we explicitly instructed the model that once a substrate is released, USP1/UAF1 will not immediately bind and cleave that same substrate again. Released PCNA-Ub is absorbed in the general pool of available substrates and is treated equal to other ubiquitinated PCNAs present. Finally, we decided to fix the actual cleavage rate (*k_cut_*) to be equal for all cleavage steps as USP1 is always cleaving an isopeptide bond.

The full final model as written in KinTek is shown in Supplementary table 1 and a more schematic representation is shown in Figure 2B. While our aim was to write separate kinetic models for K63- and K48-PCNA-Ub_2_, we found that the same final model could be applied to both. This means that USP1/UAF1 uses a similar approach cleaving these different chains on PCNA and as such, the following paragraphs regarding the kinetic model can be applied to either chain type.

### USP1/UAF1 prefers to cleave ubiquitin-ubiquitin bond over PCNA-ubiquitin bond

In the schematic kinetic model (Fig. 2B,D) we show the path taken by a single entity of USP1/UAF1, as it cleaves PCNA-Ub_2_. Although the model provides the option for both distributive cleavage (*k_2_*) and processive cleavage (*k_4_*), USP1/UAF1 has a 100-300 to 1 preference (*k_2_:* 932 ± 554 s^-1^, *k_4_:* 5.5 ± 3 s^-1^) for distributive cleavage and thus processive cleavage is extremely unlikely to occur. USP1/UAF1 therefore starts by cleaving the distal ubiquitin from PCNA-Ub_2_, then releases PCNA-Ub first, followed by release of the distal ubiquitin. The same USP1/UAF1 will prefer to engage the next PCNA-Ub_2_ instead of PCNA-Ub, as affinity derived from the off-rate shows that USP1/UAF1’s affinity for K63-PCNA-Ub_2_ (*k_1_*: 1244 s^-1^) is 2.5- fold higher than the affinity for PCNA-Ub (*k_5_*: 2812 s^-1^). Only when the pool of available PCNA-Ub_2_ depletes and the relative availability of PCNA-Ub increases, USP1/UAF1 will start cleaving PCNA-Ub as well. After cleaving PCNA-Ub, USP1 first releases free PCNA after which it releases the proximal ubiquitin.

To generate a visual representation of this process, we used the kinetic model to simulate how the amounts of substrate and products vary over time (Fig. 2C). While chains on PCNA disappear rapidly, the amount of monoubiquitinated PCNA only declines when chains are depleted. This corresponds to our DUB assays (Fig. 1), where we observe how PCNA-Ub levels increase first and only decrease at a later stage. It demonstrates that the preference of USP1/UAF1 for cleaving PCNA-Ub_2_ can result in temporal enrichment of PCNA-Ub, which in DNA damage tolerance would promote TLS (Fig. 2E). Although the affinity for K48-PCNA-Ub_2_ is lower than for K63-PCNA-Ub_2_, as seen by its *k_1_* value, the simulations show that the temporal enrichtment of PCNA-Ub can also take place for K48-PCNA-Ub_2_.

The kinetic model also allows calculation of detailed reaction parameters (Fig. S4C,D) that allows us to calculate the catalytic efficiency (k_cat_/k_m_) of the reaction (Table 1). We use Michaelis-Menten analysis for PCNA-Ub and distally labeled PCNA-Ub_2_. For the proximally labeled PCNA-Ub_2_ substrates however, Michaelis-Menten kinetics are not appropriate. USP1/UAF1’s preference for cleaving the distal ubiquitin causes a delay in cleavage of the proximal TMR-labeled ubiquitin, as shown in the kinetic simulation for PCNA-Ub^TMR^-Ub (Fig. S4E,F). We therefore use this substrate inhibition model to calculate the reaction rates (Fig. S4C,D) for proximally labeled PCNA-Ub_2_ instead of the Michaelis-Menten model.

**Table 1.**
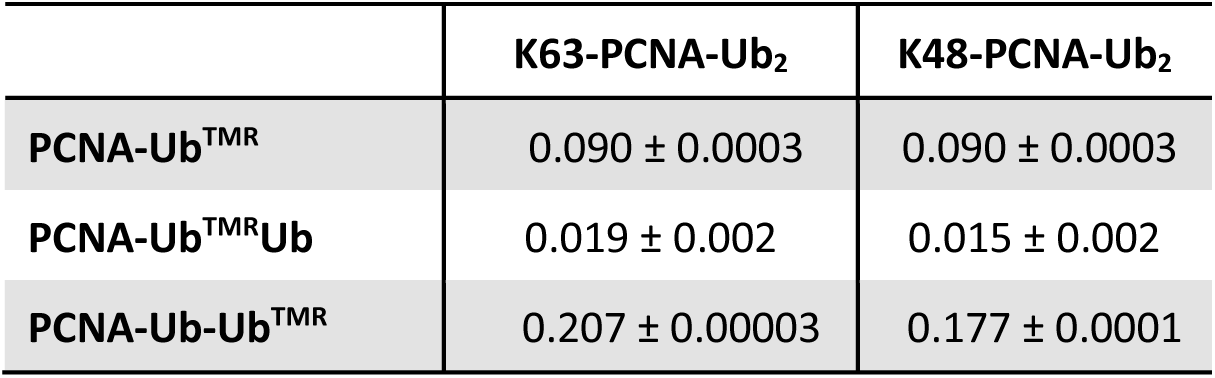
Catalytic efficiencies (K_cat_/K_M_) of USP1/UAF1 on K63-PCNA-Ub_2_, K48-PCNA-Ub_2_ and PCNA-Ub. Catalytic efficiencies (µM^-1^s^-1^) derived from Kintek model (*Fig. 2C,D, Table S1*). Full analysis is shown in *Supplementary figure S4C,D*.

Similar to the kinetic model itself, the catalytic efficiencies (K_cat_/K_m_) of USP1/UAF1 for distally labeled K63- and K48-PCNA-Ub_2_ are comparable. These catalytic efficiencies show that USP1/UAF1 has a ∼10 fold preference for cleaving Ub-Ub bonds (either 0.207 nM^-1^s^-1^ or 0.156 nM^-1^s^-1^ for K63-PCNA-Ub-Ub^TMR^ or K48-PCNA-Ub-Ub^TMR^ respectively) over the PCNA-Ub bond in PCNA-Ub_2_, (0.019 nM^-1^s^-1^or 0.015 nM^-1^s^-1^ for K63-PCNA-Ub^TMR^-Ub or K48-PCNA-Ub^TMR^-Ub respectively). The low catalytic efficiencies for proximally labeled PCNA reflect USP1/UAF1’s exo-cleavage and bind and release mechanism, as the enzyme would need to cleave the distal ubiquitin before we could detect the release of the proximal Ub^TMR^ (Fig. 2B,C, Fig S2E,F), resulting in these lower catalytic efficiencies. Cleavage of monoubiquitinated PCNA itself (PCNA-Ub: 0.090 nM^-1^s^-1^) is also less efficient compared with exo-cleavage of the distal ubiquitin.

### Structure of USP1/UAF1 in complex with K63-linked diubiquitin

To gain structural insights into USP1/UAF1’s mechanism on ubiquitin chains and study why it prefers exo-cleavage, we performed cryo-EM single particle analysis on full length USP1/UAF1 crosslinked with the K63-linked diubiquitin VA probe warhead (Ub_VA_Ub). We obtained a reconstruction of the complex at 3.0 Å resolution (Fig. 3).

**Figure 3.**
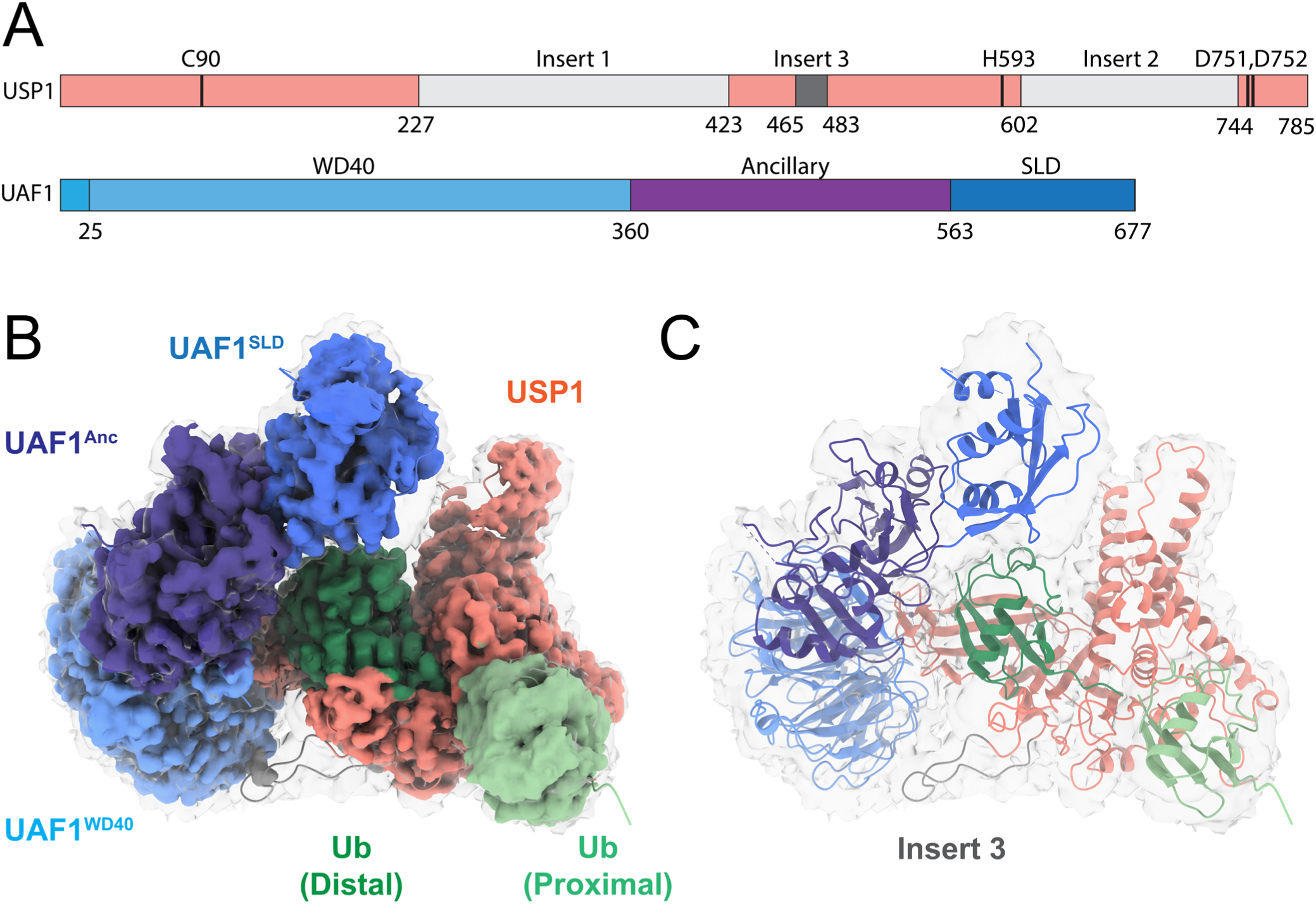
Cryo-EM structure of USP1/UAF1 bound to K63-Ub_VA_Ub. **A)** Domain architecture of USP1 and UAF1. Density was resolved for USP1’s insert 3 (dark grey), but not for insert 1 and insert 2 (light grey). **B)** Final map of 3.0 Å final structure. DeepEMhancer sharpened map is coloured as in A and shown at threshold 0.001. The grey transparent map is the non-sharpened map and shown at threshold 0.015 to accentuate density for flexible regions. **C)** Cryo-EM structure of USP1/UAF1 bound to K63-linked Ub_VA_Ub probe, showing atomic model in the non-sharpened final density map.

The overall structure of USP1/UAF1 is in concordance with existing USP1/UAF1 structures (7AY0, 7AY1, 7AY2 ((Rennie et al., 2021)). USP1 interacts with UAF1 primarily via the UAF1 β-propeller domain, through the tip of its fingers sub-domain, as seen in previous structures of USP1 and consistent with USP12 and USP46 complexes with UAF1 (Dharadhar et al., 2016; Li et al., 2016; Yin et al., 2015; Zhu et al., 2019).

The structure reveals how USP1/UAF1 binds the K63-linked diubiquitin chain. As expected for an exo-cleavage mechanism, the distal ubiquitin is placed in the USP1 hand, positioning its ubiquitin tail for catalysis in the active site (Fig. S6A,B). The proximal ubiquitin is positioned at the base of the USP1 thumb in what looks like a binding pocket. They are linked by the VA-type warhead that is cross-linked to the catalytic cysteine (C90) of USP1 and links the distal and proximal ubiquitin (Fig. S6B,D). In this structure, D751 is facing the catalytic histidine (H593) (Fig. S6E). We have previously shown that this aspartate is dispensable for catalysis and that D752 instead is critical for catalysis (Keijzer et al., 2024). We built D752 to face the catalytic histidine, which follows the canonical positioning of this residue. However, there is also density to accommodate an alternative rotameric state, where D752 faces up towards N85 (Fig. S6E). The alternative state of D752 was observed in one earlier structure (7AY1 (Rennie et al., 2021)), but not in others (Rennie et al., 2021, 2024).

An unexpected feature in this structure is that insert 3 is partially ordered (residues 465 – 483), in contrast to previous structures of USP1/UAF1 (7AY0, 7AY1, 7AY2), (Rennie et al., 2021) and creates an additional interaction with the WD40 β-propeller domain of UAF1 (Fig. S6C). The interaction is facilitated through three hydrogen bonds: USP1^E473^ with UAF1^S165^ and UAF1^K203^ and UAF1^K168^ with the main chain of USP1^S475^.

### UAF1 plays a minor role in exo-cleavage preference for K63-PCNA-UbN

The structure suggests that positioning of UAF1 (Fig. 4A), may prevent cleavage via base- or endo-cleavage, especially the SLD and ancillary domain. These domains may need to move to accommodate distal ubiquitins when endo- or base-cleaving ubiquitin chains. To test if these domains would affect USP1/UAF1’s cleavage mechanism, we generated and purified deletion constructs and analyzed activity of USP1/UAF1^ΔSLD^ (res. 9 – 563) and USP1/UAF1^ΔAnc-SLD^ (res. 25-360) (Fig. 4B) on K63-PCNA-Ub_N_ and K48-PCNA-Ub_N_. These deletions did not have a major effect on the thermal stability of the protein (Fig. S7). Deletion of both the SLD and Ancillary domain together causes a slight decrease in activity and causes a more flexible mixture between exo- and endo-cleavage. Monoubiquitinated PCNA still accumulates, showing that USP1/UAF1^ΔAnc-SLD^ retains its preference for the Ub-Ub bond. USP1/UAF1 ^ΔSLD^ on the other hand behaves like wild-type USP1/UAF1 in both cleavage pattern preference and activity. Thus, despite opening up more space to accommodate ubiquitin chain binding for endo- or base- cleavage, these USP1/UAF1 variants mostly retain their preference for exo-cleavage.

**Figure 4.**
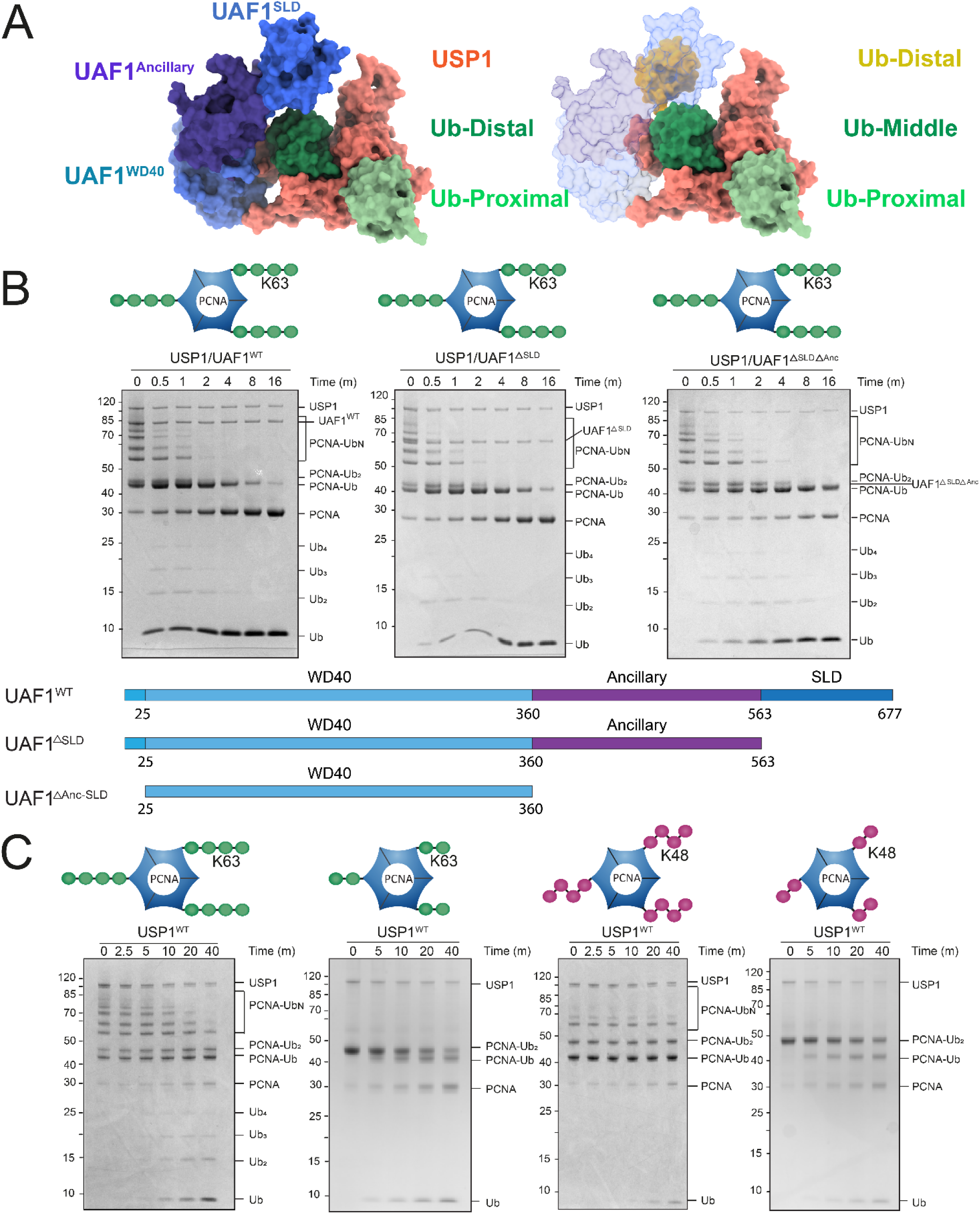
Analysis of the role of steric hindrance by UAF1 in exo-cleavage preference. **A)** Left: USP1/UAF1 showing the subdomains of UAF1 (WD40, Ancillary domain, SLD) in different tints of blue. Right: Theoretical depiction of USP1/UAF1 binding the chain in an endo-cleavage mechanism shows how UAF1 SLD and Ancillary domains might obstruct binding. **B)** DUB assays on K63-PCNA-Ub_N_ shows that deletion variants (USP1/UAF1 ^ΔSLD^ and USP1/UAF1^ΔSLD+ΔAncillary^) of UAF1, employ the same cleavage mechanism as full length USP1/UAF1. Constructs used for deletion mutants are shown in lower panel. **C)** DUB assays of USP1 alone on K63-PCNA-Ub_N_ and K63-PCNA-Ub_2_ (left) show that USP1 employs a mix of exo- and endo-cleavage, but does not seem to cleave at the base as indicated by cleavage of K63-PCNA-Ub_2_. On K48-PCNA-Ub_N_ and K48-PCNA-Ub_2_ (right panel) USP1’s activity is even more reduced compared to K63-PCNA-Ub_N_, but still uses exo-cleavage.

We then tested activity of USP1 without UAF1, to understand their relative contribution to the mechanism. These experiments required higher concentrations of USP1, as the enzyme alone is less active (Cohn et al., 2007, 2009). When cleaving K63-PCNA-Ub_N_, USP1^WT^ generates an equal mixture of free chains and free ubiquitin at t = 5 minutes. Analysis of USP1^WT^ on K63-PCNA-Ub_2_ confirms that USP1 still opts for the Ub-Ub bond rather than the PCNA-Ub bond. USP1^WT^ therefore uses a mix of exo- and endo-cleavage on K63-PCNA-Ub_N_ (Fig. 4C), but still does not opt for base-cleavage. For K48 chains USP1^WT^ is hardly active and can only be seen to generate free ubiquitin. In line with previous findings (Dharadhar et al., 2021), the USP1 inserts may obstruct binding and activity on K48-chains without UAF1.

These results show that some steric hindrance might be caused by UAF1 or UAF1’s Ancillary and SLD domain together, but that the preference for the Ub-Ub bond is caused by USP1 alone. The results also show that the UAF1 WD40-domain is sufficient for activation of USP1, indicating that it is sufficient to resolve the autoinhibition from insert 1 and insert 3. The observed interaction of UAF1^WD40^ with insert 3 (Fig. S6C), could contribute to this. UAF1’s SLD domain alone does not appear to play a role in PCNA-Ub or PCNA-Ub_N_ deubiquitination, in contrast to its role for FANCI/FANCD2 deubiquitination (K. Yang et al., 2011).

### Hyperactivation of USP1 by mutating a conserved negative patch

The structure shows how the proximal ubiquitin is positioned in a pocket next to the USP1 thumb, which consists mostly of negatively charged residues (Fig. 5A,B,C). Two of these residues (D751 and D752) are well conserved active site residues and E754, which faces the proximal ubiquitin, is also well conserved. The two remaining residues in this patch (D172 and E173) are positioned in a different loop. These residues are well conserved in USP1 homologs in evolution (Fig. 5D), but are not conserved in USP paralogs, except for USP30 (Fig. S8A,B).

**Figure 5.**
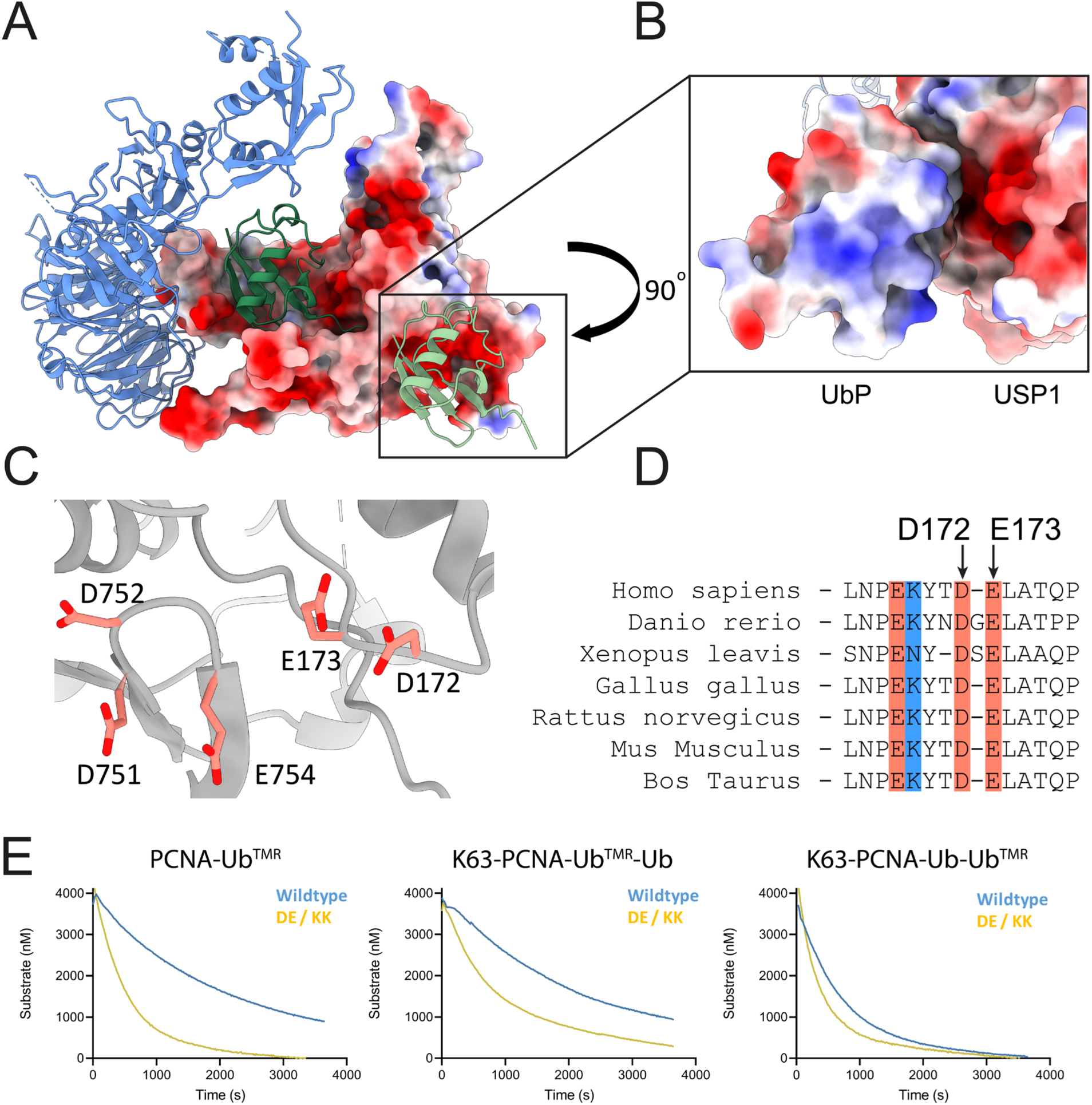
Hyperactivation of USP1 through mutations in its conserved negative patch. **A)** Electrostatic surface of USP1 indicates negatively charged binding pocket for the proximal ubiquitin. **B)** Electrostatic depiction of interface between USP1 and the proximal ubiquitin. **C)** Details of the negatively charged patch in USP1, D751, D752 are active site residues, and E754 supports their positioning. D172, E173 are on a separate loop. **D)** Sequence alignment of USP1 orthologs shows that D172 and E173 are well conserved in USP1 homologs alone. Residues are colored by charge. E) Comparison of activity of D172/E173 to lysine (DE/KK) mutant to wildtype USP1 on PCNA-Ub, K63-PCNA-Ub^TMR^-Ub and K63-PCNA-Ub-Ub^TMR^. USP1^DE/KK^/UAF1 displays a major increase in activity on PCNA-Ub bonds (read out in PCNA-Ub^TMR^ and K63-PCNA-Ub^TMR^-Ub) but only a minor increase on Ub-Ub bonds (readout in K63-PCNA-Ub-Ub^TMR^).

Only four USP structures complexed with Ub_2_ are available for comparison (USP21 + M1-Ub_2_; 2Y5B (Ye et al., 2011), USP30 + K6-Ub_2_; 5OHP (Gersch et al., 2017), CYLD K63-Ub_2_; 3WXG, CYLD + M1-Ub_2_; 3WVE (Sato et al., 2015)). USP30 however, is bound to a K6-linked diubiquitin and as such, the proximal ubiquitin is positioned differently with regards to the catalytic domain and the residues corresponding to D172 and E173 are not involved (Fig. S8A,B). The catalytic domain of CYLD adopts a fold slightly different from the canonical USP catalytic domain fold. The proximal ubiquitin of M1 – and K63-Ub_2_ is twisted and positioned differently from the proximal ubiquitin bound to USP1, and CYLD has a protruding B-sheet which interacts with the proximal ubiquitin, which is not found in USP1 (Sato et al., 2015). USP21 interacts with an M1-Ub_2_ chain with a preference for base-cleavage and has the C-terminal tail of the proximal subunit positioned for cleavage, placing the distal ubiquitin beyond that, on the tip of the fingers subdomain.

To study the role of D172 and E173 in catalysis we mutated the DE to AA or KK respectively and compared their activity to USP1^WT^/UAF1. Strikingly, both mutants show gain-of function, a small increase in activity for USP1^DE/AA^/UAF1, and a more substantial increase for USP1^DE/KK^/UAF1 for cleavage of PCNA-Ub_N_ (Fig. S8D). Both mutants still primarily use exo-cleavage as evidenced by the production of free ubiquitin and the decrease in chain length of PCNA-Ub_N_, similar to USP1^WT^/UAF1.

We tested the mutants on fluorescently labeled K63-PCNA-Ub_2_ substrates to identify on which bond of the ubiquitin chains the reaction is enhanced. Hyper-activation of USP1^DE/KK^/UAF1 occurs on PCNA-Ub^TMR^, K63-PCNA-Ub^TMR^-Ub and only to a minor extent on K63-PCNA-Ub-Ub^TMR^ (Fig. 5E). Interestingly, this increase in activity is especially pronounced when cleaving proximally labeled K63-PCNA-Ub^TMR^-Ub and PCNA-Ub^TMR^, indicating that it is specific for PCNA-Ub bonds.

We wondered why evolution maintains the slower D172/E173 in USP1^WT^, and not the increased activity of the positively charged K172/K173. We therefore tested thermal stability of these mutants and surprisingly, USP1^DE/KK^/UAF1 shows a 5°C increase in thermal stability over USP1^WT^/UAF1 (Table S3, Fig. S8E). As stability was evidently not the evolutionary push that favors USP1^WT^ over USP1^DE/KK^, these results suggest that there is a different evolutionary advantage to retain a suboptimal USP1 that has lowered activity for monoubiquitinated PCNA. The lower activity of USP1^WT^ allows for more enrichment of PCNA-Ub and opportunities for fine tuning the DNA damage tolerance pathways.

## Discussion

In this research, we reveal that USP1/UAF1’s kinetics and cleavage mechanism on polyubiquitinated PCNA allow for enrichment of monoubiquitinated PCNA, thereby directing DNA damage tolerance. This is possible because USP1/UAF1 prefers exo-cleavage and therefore trims chains down, but also because it has a kinetic preference for Ub-Ub bonds over Ub-PCNA bonds. Our kinetic modeling shows that USP1/UAF1 prefers cleaving polyubiquitinated over monoubiquitinated PCNA for both K63- and K48-polyubiquitinated PCNA. USP1/UAF1 uses a bind, cleave and release mechanism, releases PCNA-Ub and moves to cleave a different polyubiquitinated PCNA, causing PCNA-Ub to accumulate.

This kinetic preference gives USP1/UAF1 multiple roles (Fig. 6). In the absence of damage, when there is little PCNA ubiquitination, it will effectively remove monoubiquitin, preventing unscheduled onset of DDT. Upon fork stalling, when both mono- and polyubiquitination take place, it will focus on trimming of chains, and temporarily enrich for monoubiquitinated PCNA. This will protect against formation of chains on PCNA, thereby preventing unscheduled HR-mediated bypass or PCNA degradation and promoting TLS. If a lesion proves too challenging for TLS or if bypass takes too long, sufficient K63-linked chains could form, resulting in HR-mediated DDT. Since USP1 is highly active, removal of USP1 through its autocleavage mechanism, which occurs e.g. upon exposure to UV-light (Huang et al., 2006) might be required to allow DDT mechanisms to proceed.

**Figure 6.**
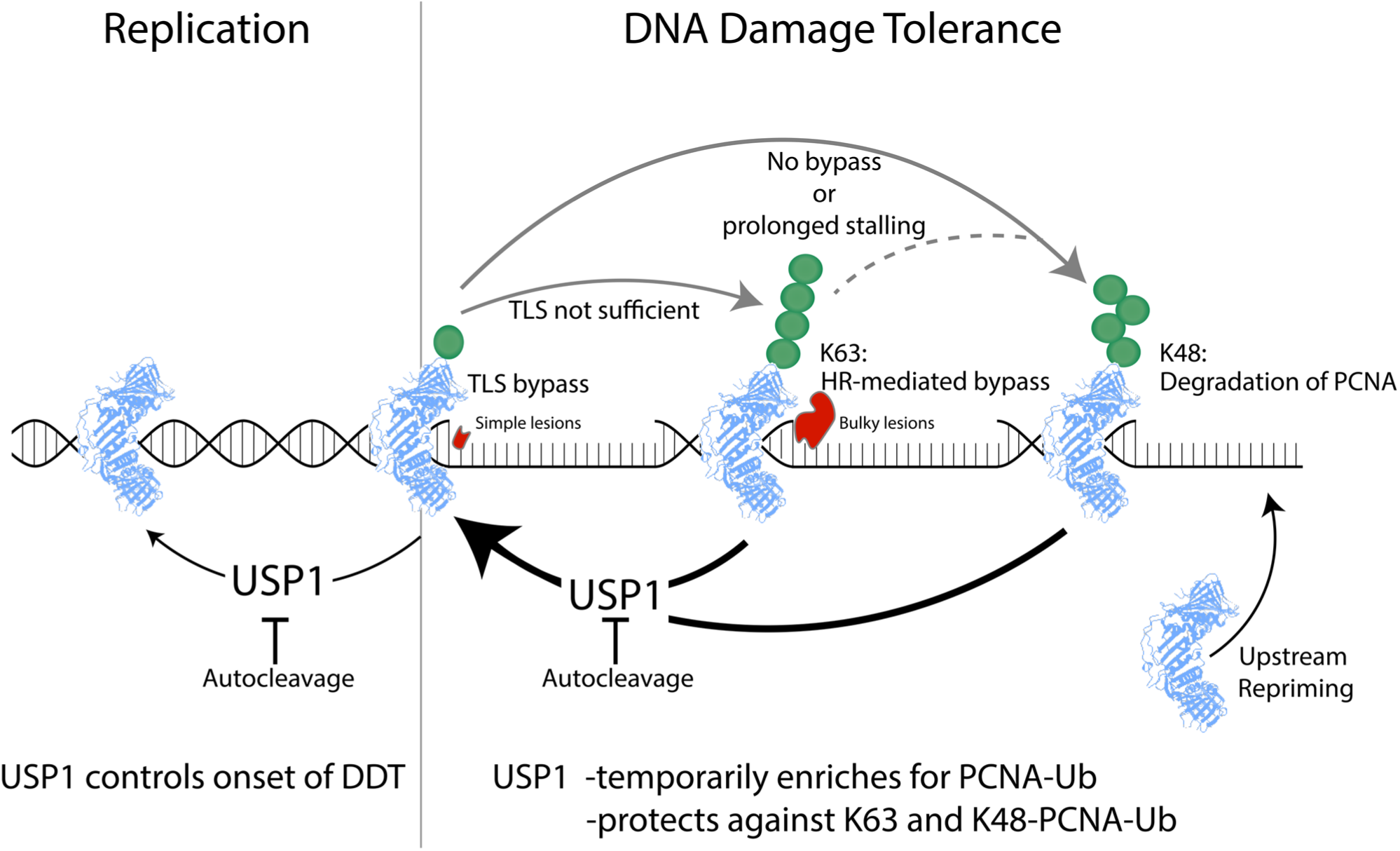
Role of USP1/UAF1’s kinetic preference in DNA damage tolerance. In a normal replicating fork, USP1 will prevent premature monoubiquitination. Upon stalling at a DNA lesion, PCNA is monoubiquitinated at lysine 164, which promotes TLS, offering rapid bypass of small and simple DNA lesions. Upon successful TLS bypass, USP1/UAF1 deubiquitinates PCNA which allows PCNA to switch back to replication. When DNA lesions are too bulky or complicated and cannot be bypassed by TLS, K63-PCNA-Ub_N_ is produced, which promotes HR-mediated bypass. As HR-mediated bypass is most likely slower and more complex compared with TLS and it is prevented when TLS is sufficient. USP1 initially safeguards against undesired K63-ubiquitination by processively cleaving the K63-PCNA-Ub_N_, thereby reverting back to PCNA-Ub allowing for TLS. Prolonged stalling will eventually result in K63-linked chain formation and autocleavage of USP1 can accelerate this. When lesions prove to be too complicated for either type of bypass, or when another factor causes prolonged stalling of PCNA, PCNA-Ub is K48-poly-ubiquitinated and subsequently degraded. Upon degradation of PCNA, repriming allows a new PCNA to be loaded upstream. USP1 protects PCNA against undesired degradation and turns to TLS before a return to replication, thereby protecting replication and cell survivability.

Only when replication impediments cause severe replication stress and cannot be resolved by either TLS or HR-like bypass, K48-ubiquitination and subsequent removal of PCNA would offer a solution. Followed by upstream PrimPol mediated repriming (Bainbridge et al., 2021) replication would still be able to continue, but would leave behind a ssDNA gap. In turn, these ssDNA gaps can be filled post replicative in a PCNA ubiquitination independent manner (Nusawardhana et al., 2024).

This DNA damage tolerance pathway choice is affected by many factors. It will certainly be regulated by local concentrations of E3 ligases. Although USP1/UAF1 is efficient at deubiquitinating PCNA, its affinity for PCNA is relatively low (Dharadhar et al., 2021). Competition with the different relevant E3 ligases could shift the balance in DDT. Alternatively, the type of DNA damage could decide which pathway is required. TLS may rapidly bypass smaller DNA lesions, whereas bulky or complicated lesions, that do not fit in the TLS polymerase active site would require HR-mediated bypass. Additionally, since specific DNA lesions require specific TLS polymerases for bypass (Friedberg et al., 2005), it is possible that HR-mediated bypass is important when the relevant TLS polymerase is not available.

Our findings of USP1/UAF1 as a kinetic regulator of DDT pathway choice is supported by the evolutionary conservation of two negatively charged residues (D172, E173) in USP1. Mutating these residues to positively charged lysines, generates a hyperactive USP1 which is enhanced for cleavage of the PCNA-Ub bond. Conservation of the sub-optimal situation suggests that this lowered activity is vital to USP1s role in fine-tuning DDT pathways.

Our structure also revealed a partially ordered insert 3 (residues 465 – 483), which together with insert 1 (residues 227 – 423), auto-inhibits USP1 (Dharadhar et al., 2021). UAF1 binding resolves the autoinhibition, and therefore it is interesting to note that insert 3 anchors at the UAF1 WD40 domain. One of the USP1 residues in that interface, S475 frequently phosphorylated (Hornbeck et al., 2015) and there is cell-based evidence that Ser475 is important for USP1/UAF1 activity (Olazabal-Herrero et al., 2016). It is therefore possible that insert 3 anchoring to UAF1, is enhanced or reoriented by USP1^S475^ phosphorylation. This binding of insert 3 to UAF1 could prevent collapse of insert 3 on blocking loop 1 and 2 (BL1 residues 536 -551, BL2 residues 564 – 592) in the USP1 catalytic domain.

The importance of D752 in catalysis (Keijzer et al., 2024) remains interesting, and our cryo-EM density map is ambiguous on its rotameric state, making it hard to interpret its role. It is possible that D752 is indeed flexible and fulfills a dual role. Facing the active site, D752 could support the other catalytic residues whereas its alternative conformation could reinforce the oxyanion hole stabilizing role of N85. However, since the active site is currently engaged with the VA-warhead, it is possible that the active site is trapped in an intermediate state and that this affects positioning of D752.

Despite their different spatial organization, USP1/UAF1 uses the same cleavage mechanism on both K63 and K48 chains. While processing the cryo-EM dataset, resolving density of the proximal ubiquitin proved difficult at first and required extensive processing. This suggests that there is some flexibility in the position of the proximal ubiquitin. This minor flexibility could explain why USP1 is able to occasionally perform endo-cleavage on K63-PCNA-Ub_N_. K48-chains on the other hand adopt a more rigid conformation than K63-chains, which could explain why USP1/UAF1 is only able to use exo-cleavage on K48 polyubiquitinated PCNA.

The findings in this study reveal an unexpected role of USP1/UAF1, where intrinsic kinetic mechanisms play a role in regulating DNA damage tolerance. With USP1 inhibitors currently undergoing clinical trials, understanding the different functions of USP1 becomes increasingly relevant. This study yields novel insights into regulation of DNA damage tolerance and helps us understand more about the role of USP1 in regulating ubiquitinated PCNA, by protecting against K63- or K48-PCNA-Ub_N_ and promoting mono over polyubiquitination.

## Acknowledgements

We thank personnel of NeCEN and Xiaohu Guo from the NKI cryo-EM facility for assistance in data collection. Thanks to Andrea Murachelli and Shun Hsiao Lee with cryo-EM data processing and Robbie Joosten for helping with model building. The authors would like to acknowledge the Research High Performance Computing (RHPC) facility of the Netherlands Cancer Institute (NKI) which was used for cryo-EM data processing. This work is part of the Oncode Institute, which is partly financed by the Dutch Cancer Society. Research at the Netherlands Cancer Institute is supported by institutional grants of the Dutch Cancer Society and the Dutch Ministry of Health, Welfare and Sport. This study was supported by NWO grants OCENW.KLEIN.131, TOP714.016.002 and Health Holland grant LSHM21048-H045.

## Data availability

Cryo-EM reconstruction and structure coordinates are available on the Electron Microscopy Database (EMDB) and to the PDB under the following accession code: PDB: 9HNW, EMD: 52316.

## Methods

### Constructs and mutagenesis

USP1 (pFastbac-HTb, N-terminal His-tag, res. 21-785, G670A + G671A (Huang et al., 2006)) variants (USP1^D172A/E173A^, USP1^D172K/E173K^) were generated using site-directed mutagenesis. UAF1 (pFastbac1, N-terminal strep-tag, res. 9-677) and its deletion variants (UAF1^ΔSLD^ (res. 9 – 563), UAF1^ΔAnc-SLD^ (res. 25-360) were generated using 5’-phosphorylated primers and site-directed mutagenesis. USP1 and UAF1 expressed and purified for cryo-EM had short C-terminal extensions as described previously (Keijzer et al., 2024). Enzymes used for *in vitro* ubiquitination assays were expressed using the following constructs: hUBA1 (pET3a-UBA1), UbCH5C^S22R^ (pET28a-UbCH5C^S22R^), Ubc13/Mms2 (pGEX-UBC13, pET16b-MMS2), UBE2K (pETM30-GST-UBE2K^1–155^) and RNF8^RING^ (pGEX6p1-RNF8^RING(351-485)^).

### Protein purification

USP1/UAF1 was copurified by coexpression of USP1 and UAF1 in Sf9 cells, using a protocol adapted from earlier work (Dharadhar et al., 2021). After lysis and centrifugation, we first performed step-affinity purification using streptactin-XT beads (IBA), followed by nickel-affinity and size-exclusion chromatography, which improved our final yield of purified protein. PCNA, ubiquitin and enzymes for *in vitro* ubiquitination were expressed and purified following previously described methods: UBA1 (Hibbert et al., 2011), UbCH5C^S22R^ (Brzovic et al., 2006) and PCNA were purified following (Hibbert et al., 2011; Hibbert & Sixma, 2012). RNF8^RING^ following (Mattiroli et al., 2012). UBC13/MMS2 following (Pickart & Raasi, 2005), UBE2K^1-155^ following (Pichler et al., 2005). Ubiquitin variants (Ub^A28C^, Ub^K63R^, Ub^K48R^, Ub^A28C/K63R^, Ub^A28C/K48R^) were purified following (Hibbert & Sixma, 2012).

### Purification of PCNA-Ub, PCNA-Ub_2_, PCNA-Ub_3_ and PCNA-Ub_N_ variants

Different variants of mono- or diubiquitinated PCNA were made using either unlabeled ubiquitin (Ub) or TAMRA (TMR)-labeled ubiquitin (Ub^TMR^). For fluorescent labelling we made use of Ub^A28C^, to generate three different types of TMR-labeled ubiquitin variants (Ub^TMR^, Ub^K63R-TMR^, Ub^K48R-TMR^). The labelling procedure followed previously described methods (Dharadhar et al., 2019).

Monoubiquitinated PCNA (PCNA-Ub and PCNA-Ub^TMR^) was made enzymatically using UBA1 and UbCH5C^S22R^ (UBE2D3) (Hibbert & Sixma, 2012), aiming to monoubiquitinate every monomer of PCNA. To generate the K63-PCNA-Ub_VA_Ub activity-based probe, the protocol was adapted, substituting K63-linked Ub_VA_Ub for Ub ((Mulder et al., 2014) UbiQ-087 UbiQ Bio B.V.), and performing the reaction at 4°C for 45 minutes, so that on average only one of the monomers is modified with Ub_VA_Ub. Note that in Mulder et al., 2014, Ub_VA_Ub is termed diUb VME.

For producing K63-linked PCNA-Ub_2_ variants (K63-PCNA-Ub-Ub, K63-PCNA-Ub^TMR^-Ub, K63-PCNA-Ub-Ub^TMR^), 12 μM PCNA-Ub or PCNA-Ub^TMR^ was mixed with 32 μM Ub^K63R^ or Ub^K63R-TMR^, 200 nM UBA1, 4 μM UBC13/MMS2 and 2 μM RNF8^RING^ in reaction buffer (50 mM HEPES pH 7.5, 150 mM NaCl, 1 mM DTT, 5mM MgCl2, 5mM ATP). Diubiquitination was performed for 60 minutes at 37 °C to ensure that every monomer was diubiquitinated, after which PCNA-Ub-Ub^K63R^ was purified using anion-exchange and size exclusion as described for PCNA-Ub (Hibbert & Sixma, 2012). PCNA-Ub_VA_Ub-Ub^TMR^, was produced using the same protocol, adding Ub^K63R-TMR^ to K63-PCNA-Ub_VA_Ub.

K48-linked PCNA-Ub_2_ variants (K48-PCNA-Ub-Ub, K48-PCNA-Ub^TMR^-Ub, K48-PCNA-Ub-Ub^TMR^) were produced in an analogous manner: 16 μM PCNA-Ub or PCNA-Ub^TMR^ was mixed with 100 μM Ub^K48R^ or TMR-labeled Ub^K48R^ and 20 μM E2-25K^1-155^, 10 μM RNF8^RING^ for 60 minutes while shaking at 500 rpm at 37°C. The sample was purified following the steps described for other ubiquitinated PCNA variants. K63-PCNA-Ub_N_ and K48-PCNA-Ub_N_ were generated using the same methods as their PCNA-Ub_2_ counterparts, using 100 μM WT ubiquitin.

### PCNA-Ub, PCNA-Ub_2_ and PCNA-Ub_N_ deubiquitination assays by SDS page analysis

DUB activity assays were performed at RT, unless stated otherwise, in reaction buffer consisting of 20 mM Hepes, pH 7.5, 150 mM NaCl, and 2 mM DTT. 200 nM USP1/UAF1 was prepared at a 2x concentration (400nM USP1/UAF1). Different ubiquitinated PCNA variants (0.67 μM of PCNA-Ub, 0.67 μM PCNA-Ub_2_ or 1.33 μM PCNA-Ub_N_ (trimer concentration) were prepared at 2x concentration as well (1.33 μM for PCNA-Ub/PCNA-Ub_2_ and 2.67 μM for PCNA-Ub_N_). Before mixing, 7.5 μl substrate and 7.5 μl enzyme were separately added to 5 μl 4x SDS-page loading buffer as t_0_. Reaction was initiated by mixing remaining enzyme and substrate 1:1, after which 15 μl samples were taken at the indicated time points. Samples were loaded on a NuPAGE 4-12% Bis-Tris SDS gel and bands were separated by running for 30 min at 160 V. Whenever TMR-labeled substrates were used, gels were first imaged using a Typhoon FLA 9000 biomolecular gel scanner (Cytiva) to visualize the TMR signal, after which the gel was stained with Coomassie. Coomassie blue stained gels were imaged using a Gel Doc EZ imaging system (Bio-Rad Laboratories, Inc.). For assays testing USP1 without UAF1 the same protocol was followed, except the reaction was performed at 37°C with final concentrations of 1 µM USP1, 6 µM PCNA-Ub_N_ or 4 µM PCNA-Ub_2_, and 5 µL timepoints were taken as indicated.

### K63-Ub_VA_Ub conjugation assays

Ubiquitin probe crosslinking analysis was performed by mixing enzyme and substrate to final concentrations of 200 nM USP1/UAF1 and 2 μM PCNA-Ub_VA_Ub-Ub^TMR^ (trimer), PCNA-Ub_VA_Ub (trimer) or 6 µM Ub_VA_Ub. Enzyme and substrate were prepared at 2x concentration in reaction buffer (20 mM Hepes pH 7.5, 150 mM NaCl, and 2 mM DTT). Before mixing, 7.5 μl substrate and 7.5 μl enzyme were separately added to 5 μL 4x SDS-page loading buffer as t_0_. 15 μl samples were taken at the indicated time points. Samples were loaded on a NuPAGE 4-12 % Bis-Tris SDS gel, and were separated by running for 30 min at 160 V. The gel containing PCNA-Ub_VA_Ub-Ub^TMR^ was imaged for fluorescent TMR signal using a Typhoon FLA 9000 biomolecular imager (Cytiva), after which all gels were stained using Coomassie.

### Fluorescence Polarization activity assays

For fluorescence polarization (FP) activity assays, TMR-labeled mono- or diubiquitinated variants of PCNA were used (PCNA-Ub^TMR^, PCNA-Ub^TMR^-Ub^K63R^, PCNA-Ub-Ub^K63R-TMR^, PCNA-Ub^TMR^-Ub^K48R^, PCNA-Ub-Ub^K48R-TMR^). All reactions were performed in a 384-well plate (Corning, flat bottom, low flange) on a PHERASTAR plate reader (BMG Labtech), using a 540/590/590 nm filter block (BMG Labtech) at 25 °C. The FP-buffer consisted of 20 mM Hepes, pH 7.5, 150 mM NaCl, 5 mM DTT and 0.05% Tween-20.

Single-point assays were performed using 10 nM USP1/UAF1 and 1 μM of ubiquitinated PCNA, prepared separately at 2x concentration in FP-buffer. 10 μl of enzyme was pipetted to the plate in triplicates. To initiate the reaction, 10 μl of substrate was pipetted to the plate, after which measurement was started immediately. Release of Ub^TMR^ could be measured by the increase in fluorescence polarization.

For the global kinetic fit analysis, 10 nM USP1/UAF1 was prepared at 2x concentration in FP-buffer. USP1/UAF1 was tested against a substrate dilution series ranging from 4 μM to 0.5 μM, prepared at a 2x concentration. 10 µl of each substrate concentration from the dilution series was pipetted to a 384-well plate in triplicate. The experiment was started by injecting 10 µl of 2x USP1/UAF1 to each well using the built-in syringe of the instrument. The first datapoint was 26 seconds after injection, with subsequent datapoints taken every 14 seconds.

### Microscale thermophoresis

For MST analysis, the USP1/UAF1 was fluorescently labelled at random sites. For this, all cysteines in USP1/UAF1 were first reduced by adding 5 mM DTT to 12 µL final volume containing 20 µM USP1^WT^/UAF1^WT^ or USP1^KK^/UAF1, followed by incubation for 10 minutes at room temperature, after which the sample was buffer exchanged to 20 mM Hepes, 150 mM NaCl using a desalting column (Zeba Spin 40K MWCO, 75µl, Thermofisher). Then 12µL USP1/UAF1 was mixed with 40 µM of fluorescent label (DY-547P1-maleimide, Dyomics) and incubated for 30 minutes at room temperature. 5 mM DTT was added to inactivate the non-ligated DY-547P1, after which the sample was buffer exchanged to 20 mM Hepes (pH 7.5), 150 mM NaCl, 2 mM DTT to get rid of non-ligated fluorescent label.

Binding analysis of monoubiquitinated PCNA made use of non-cleavable Ub^L73P^ (Békés et al., 2013). 5 nM labeled USP1/UAF1 was tested against a dilution series of 250 µM to 15.625 nM substrate (PCNA, Ub^WT^, PCNA-Ub^L73P^ (Békés et al., 2013)), each prepared at 2-fold higher concentration in 20 mM Hepes pH 7.5, 150 mM NaCl, 2 mM DTT, 0.05% Tween. 15 µl of enzyme was mixed with 15 µl of different substrate concentrations, loaded into microscale thermophoresis (MST) capillaries in duplicates and inserted into the MST system (NanoTemper). Two biological replicates were performed for each MST experiment. Data was analyzed and affinities were derived using NanoTemper’s internal software.

### Kintek/Michaelis Menten analysis

The data from the FP activity assay and MST were analysed using global kinetic modelling using KinTek Explorer Software (KinTek Corporation). A comprehensive enzymatic kinetic model was constructed to study USP1/UAF1 activity across all tested substrates. These included anisotropy readouts from FP assays for substrates PCNA-Ub^TMR^, K63-PCNA-Ub^TMR^-Ub, K63-PCNA-Ub-Ub^TMR^, K48-PCNA-Ub^TMR^-Ub, K48-PCNA-Ub-Ub^TMR^, as well as MST measurements of USP1/UAF1 binding to PCNA, PCNA, Ub^WT^ and PCNA-Ub^L73P^. Additionally, data from the enzymatic cleavage of the minimal substrate Ub^Rho^ by USP1/UAF1 (Keijzer et al., 2024) were incorporated into fitting of the model.

The initial model accounted for all conceivable reaction pathways and possible intermediates in the enzymatic process. To simplify the model, following assumptions were made: in FP activity assays, PCNA substrates were treated as monomers having a single ubiquitination site. On-rate constants *k*_on_ were fixed at diffusion-limited values (10^8^ M^-1^s^-1^). The isopeptide bond cleavage rate was unified across all reaction steps and off-rate constants (*k_2_ to k_6_*) were unified for similar substrates and products, irrespective of labelling. For instance, *k_off_* was identical for K63-PCNA-Ub-Ub^TMR^ and K63-PCNA-Ub^TMR^-Ub. All constants were shared between K63- and K48 labeled substrates except for *k_1_* which are not shared between K63-PCNA-Ub_2_ and K48-PCNA-Ub_2_. To account for potential pipetting errors during the experiment, the model incorporated a possible variation of up to 5% in the initial substrate concentrations.

After constructing the initial model, iterative fitting steps were performed. In each step, kinetic constants that had no impact on fit quality were eliminated, along with their corresponding reaction steps, reducing model complexity. This process continued until no further simplification was feasible. The resulting model captures two pathways following distal Ub cleavage: distributive and processive (Table S1, Fig. 2B). This model simulates the enzymatic cleavage of PCNA-Ub-Ub, depicting changes in levels substrate, product, and PCNA-Ub intermediate product (Fig. 2C). Furthermore, it enables simulations under Michaelis-Menten conditions for each substrate. Furthermore, the model allows for simulations under Michaelis-Menten conditions for each substrate. The initial reaction rates derived from these simulations were used to calculate the catalytic efficiency *k_cat_/K_m_* (Table 2) for each substrate using GraphPad Prism (GraphPad Software LLC). For PCNA-Ub^TMR^ and PCNA-Ub-Ub^TMR^, the calculations were based on the Michaelis-Menten model, while for PCNA-Ub^TMR^-Ub, the substrate inhibition model was applied.

### Cryo-EM sample preparation

USP1/UAF1 (final 4 µM) was conjugated to K63-linked Ub_VA_Ub (UbiQ Bio B.V.) (final 16 µM) by incubating for 30 minutes at 37 °C in reaction buffer (20 mM HEPES pH 7.5, 150 mM NaCl, 2mM TCEP) to generate 4 µM (0.75mg/ml) of complex. Grids (Cu 300 Holey Carbon R1.2/1.3 (Quantifoil Micro Tools GmbH)) were glow-discharged for 1 minute at 30 mA using a GloQube Plus (Quorum) just before use. 3 μL of sample was applied to the grid and vitrified using a Vitrobot (Mark IV chamber (Thermo Fisher Scientific)) maintained at 4 °C and 100% humidity.

### Cryo-EM data collection

Data was collected at NeCEN using a 300kV Titan Krios electron microscope (Thermo Fisher Scientific) with a K3 BioQuantum direct electron detector (Gatan). EPU-2.10.0 was used for automated data collection in super-resolution mode using AFIS centering. 5833 movies of 50 frames were acquired in super resolution mode at 0.418 Å/pixel size (Counted super-resolution) with a total dose of 50 e^−^/Å^2^ (14 e^−^/pixel/s). Defocus values ranged between -1.2 μm and -2.4 μm.

### Cryo-EM data processing

Data were imported and processed using CryoSparc (Punjani et al., 2017). Movies were binned 2x and motion corrected using patch motion correction (default settings), after which they were further processed using patch CTF estimation (default settings). An initial Topaz ((Bepler et al., 2019) deep learning particle picking model was trained using manually picked particles, picked on a 100 micrograph subset. Particles were cleaned up using 2D classification and ab initio reconstruction and were used to train a new Topaz model. This Topaz model was then applied to the full dataset. The resulting particles were cleaned up using 2D classification, ab initio reconstruction and 3D classification, followed by NU-refinement. At this point, the particles were subjected to two different methods in parallel: **1):** Iterative training of a new Topaz model and **2):** Template picking using 2D projections of the NU-refined model.

1. The NU-refined particles were used to train a new Topaz model, after which particles were cleaned up again. This process was repeated until eventually an improved Topaz model was generated. These were extracted with 4x binning (original box size: 360 pix). Particles were cleaned up using 2D classification and ab initio reconstruction, after which they were re-extracted using a 280 pixel box size. A 1 class 3D classification was run to generate a single 3D volume from all particles. This was followed by NU-refinement, global and local CTF refinement and second NU-refinement. This resulted in a 3D density map with improved density for insert 3, which was not present in the 3D volume generated using approach **2)**.
2. 2D projections of the NU-refined model were used for Cryosparc’s template picker performed on the entire dataset and were extracted using a 512 pixel box size (binned to 256 pix). 2D classification was used to remove non-particles. Particles were then cleaned up using subsequent rounds of ab initio reconstruction with three classes, each time picking the best class. The resulting class was used to train a new Topaz model and particles were extracted with 352 pixel box size (binned to 176 pix). Ab initio classes were generated, whose particles and 3D density maps were used for heterogeneous refinement, which was repeated iteratively until an improved 3D density map was generated. These particles were re-extracted in a 352 pixel box size without binning, followed by NU-refinement. This resulted in a final model with a better resolved SLD compared to approach **(1)**.

Particles of approach **1)** and **2)** were extracted at 360 pix and combined, removing only 2812 particles and therefore greatly increasing the number of particles. Since a 10 class 3D classification resulted in highly similar 3D volumes, the particles were combined into a single model using 3D classification and a mask covering the entire complex. The resulting 3D volume was NU-refined using the same mask, followed by local and global CTF refinement and another NU-refinement. This final 3D volume of combined particle stacks **(1)** and **(2)** had better SLD, insert 3 and proximal Ub density and improved local resolution. Until this point the density for the proximal ubiquitin was ambiguous and made it difficult to precisely place the proximal ubiquitin, suggesting some minor flexibility. We therefore used this refinement for 3D flex (Punjani & Fleet, 2023). This map was used for local refinement, with a mask covering the entire complex and the fulcrum coordinates set on the proximal ubiquitin (xyz: -33.8, -2.9, 57.4). We generated a tight mask on the entire complex and ran a final NU refinement, which improved the density for the proximal ubiquitin even more. This yielded the final model, which was sharpened using DeepEMhancer (Sanchez-Garcia et al., 2021).

### Model building

The final cryo-EM density and DeepEMhancer sharpened map were used to build the structure using real-space refinement in Phenix, PDB-Redo and manual jiggle fitting in Coot (Adams et al., 2010; Emsley et al., 2010; Joosten et al., 2014). We started by rigid body fitting the full length AlphaFold-2 models (Jumper et al., 2021) of USP1 (uniprot ID: O94782) and UAF1 (uniprot: Q8TAF3), from the EBI database, and two copies of the experimental structure of ubiquitin (PDB ID: 1UBQ) into our final density map using ChimeraX (Pettersen et al., 2021). The fitted model was real-space refined using Phenix (Adams et al., 2010), after which the fit was adjusted in Coot by jiggle fitting and manual rebuilding. We used Coot to generate and fit the 4-aminobutyric acid linker (termed ABU in structure) in the active site with restraints according to ABU. The model was submitted to PDB-REDO (Joosten et al., 2014), together with the restraints file for ABU, which improved the overall model. This model was examined using Coot, after which we manually corrected remaining model geometry issues, class score and rotamer outliers. This model was was submitted for one more round of Phenix real-space refinement. The output model was checked a final time in Coot, correcting for the few remaining geometry and rotamer outliers. This final model was validated using the PDB-validation tool, after which it was submitted to the PDB and EMDB. Figures including the cryo-EM model were made using USCF ChimeraX.

### Structure based sequence alignment

To compare predicted structures from all human USPs, predictions from the EBI Alphafold-2 database (Jumper et al., 2021) were used. All predicted structures were superimposed onto USP1’s Alphafold-2 structure, using the matchmaker function from UCSF Chimera (Meng et al., 2006). In turn, this superposition was used for structure-based sequence alignment, using Match -> align function in Chimera. Since the output showed that except for USP1 and USP30, this loop is not found in any of the other USPs and could therefore not be shown.

## References

Adams, P. D., Afonine, P. V., Bunkóczi, G., Chen, V. B., Davis, I. W., Echols, N., Headd, J. J., Hung, L. W., Kapral, G. J., Grosse-Kunstleve, R. W., McCoy, A. J., Moriarty, N. W., Oeffner, R., Read, R. J., Richardson, D. C., Richardson, J. S., Terwilliger, T. C., & Zwart, P. H. (2010). PHENIX: a comprehensive Python-based system for macromolecular structure solution. Acta Crystallographica. Section D, Biological Crystallography, 66(Pt 2), 213–221. 10.1107/S0907444909052925

Al-Hakim, A., Escribano-Diaz, C., Landry, M. C., Donnell, L., Panier, S., Szilard, R. K., & Durocher, D. (2010). The ubiquitous role of ubiquitin in the DNA damage response. DNA Repair, 9(12), 1229. 10.1016/J.DNAREP.2010.09.011

Bainbridge, L. J., Teague, R., & Doherty, A. J. (2021). Repriming DNA synthesis: an intrinsic restart pathway that maintains efficient genome replication. Nucleic Acids Research, 49(9), 4831–4847. 10.1093/NAR/GKAB176

Békés, M., Okamoto, K., Crist, S. B., Jones, M. J., Chapman, J. R., Brasher, B. B., Melandri, F. D., Ueberheide, B. M., LazzeriniDenchi, E., & Huang, T. T. (2013). DUB-resistant ubiquitin to survey ubiquitination switches in mammalian cells. Cell Reports, 5(3), 826. 10.1016/J.CELREP.2013.10.008

Bepler, T., Morin, A., Rapp, M., Brasch, J., Shapiro, L., Noble, A. J., & Berger, B. (2019). Positive-unlabeled convolutional neural networks for particle picking in cryo-electron micrographs. Nature Methods 2019 16:11, 16(11), 1153–1160. 10.1038/s41592-019-0575-8

Bienko, M., Green, C. M., Crosetto, N., Rudolf, F., Zapart, G., Coull, B., Kannouche, P., Wider, G., Peter, M., Lehmann, A. R., Hofmann, K., & Dikic, I. (2005). Ubiquitin-Binding Domains in Y-Family Polymerases Regulate Translesion Synthesis. Science, 310(5755), 1821–1824. 10.1126/science.1120615

Biertümpfel, C., Zhao, Y., Kondo, Y., Ramón-Maiques, S., Gregory, M., Lee, J. Y., Masutani, C., Lehmann, A. R., Hanaoka, F., & Yang, W. (2010). Structure and Mechanism of Human DNA Polymerase η. Nature, 465(7301), 1044. 10.1038/NATURE09196

Brzovic, P. S., Lissounov, A., Christensen, D. E., Hoyt, D. W., & Klevit, R. E. (2006). A UbcH5/ubiquitin noncovalent complex is required for processive BRCA1-directed ubiquitination. Molecular Cell, 21(6), 873–880. 10.1016/j.molcel.2006.02.008

Cadzow, L., Brenneman, J., Sullivan, P., Liu, H., Shenker, S., McGuire, M., Grasberger, P., Sinkevicius, K., Hafeez, N., Histen, G., Chipumuro, E., Hixon, J., Krall, E., Cogan, S., Wilt, J., Schlabach, M., Stegmeier, F., Olaharski, A., & Wylie, A. (2020). Development of KSQ-4279 as a first-in-class USP1 inhibitor for the treatment of BRCA-deficient cancers. European Journal of Cancer, 138, S52. 10.1016/s0959-8049(20)31215-6

Cadzow, L., Brenneman, J., Tobin, E., Sullivan, P., Nayak, S., Ali, J. A., Shenker, S., Griffith, J., McGuire, M., Grasberger, P., Mishina, Y., Murray, M., Dodson, A. E., Gannon, H., Krall, E., Hixon, J., Chipumuro, E., Sinkevicius, K., Gokhale, P. C., … Wylie, A. A. (2024). The USP1 Inhibitor KSQ-4279 Overcomes PARP Inhibitor Resistance in Homologous Recombination–Deficient Tumors. Cancer Research, 84(20), 3419–3434. 10.1158/0008-5472.CAN-24-0293

Chiu, R. K., Brun, J., Ramaekers, C., Theys, J., Weng, L., Lambin, P., Gray, D. A., & Wouters, B. G. (2006). Lysine 63-Polyubiquitination Guards against Translesion Synthesis–Induced Mutations. PLoS Genetics, 2(7), e116. 10.1371/journal.pgen.0020116

Cohn, M. A., Kee, Y., Haas, W., Gygi, S. P., & D’Andrea, A. D. (2009). UAF1 Is a Subunit of Multiple Deubiquitinating Enzyme Complexes. The Journal of Biological Chemistry, 284(8), 5343. 10.1074/JBC.M808430200

Cohn, M. A., Kowal, P., Yang, K., Haas, W., Huang, T. T., Gygi, S. P., & D’Andrea, A. D. (2007). A UAF1-Containing Multisubunit Protein Complex Regulates the Fanconi Anemia Pathway. Molecular Cell, 28(5), 786–797. 10.1016/j.molcel.2007.09.031

Dharadhar, S., Clerici, M., van Dijk, W. J., Fish, A., & Sixma, T. K. (2016). A conserved two-step binding for the UAF1 regulator to the USP12 deubiquitinating enzyme. Journal of Structural Biology, 196(3), 437–447. 10.1016/j.jsb.2016.09.011

Dharadhar, S., Dijk, W. J., Scheffers, S., Fish, A., & Sixma, T. K. (2021a). Insert L1 is a central hub for allosteric regulation of USP1 activity. EMBO Reports, 22(4). 10.15252/embr.202051749

Dharadhar, S., Dijk, W. J., Scheffers, S., Fish, A., & Sixma, T. K. (2021b). Insert L1 is a central hub for allosteric regulation of USP1 activity. EMBO Reports, 22(4). 10.15252/embr.202051749

Dharadhar, S., Kim, R. Q., Uckelmann, M., & Sixma, T. K. (2019). Quantitative analysis of USP activity in vitro. Methods in Enzymology, 618, 281–319. 10.1016/BS.MIE.2018.12.023

Emsley, P., Lohkamp, B., Scott, W. G., & Cowtan, K. (2010). Features and development of Coot. Acta Crystallographica. Section D, Biological Crystallography, 66(Pt 4), 486–501. 10.1107/S0907444910007493

Faesen, A. C., Luna-Vargas, M. P. A., Geurink, P. P., Clerici, M., Merkx, R., Van Dijk, W. J., Hameed, D. S., El Oualid, F., Ovaa, H., & Sixma, T. K. (2011). The differential modulation of USP activity by internal regulatory domains, interactors and eight ubiquitin chain types. Chemistry and Biology, 18(12), 1550–1561. 10.1016/j.chembiol.2011.10.017

Friedberg, E. C., Lehmann, A. R., & Fuchs, R. P. P. (2005). Trading Places: How Do DNA Polymerases Switch during Translesion DNA Synthesis? Molecular Cell, 18(5), 499–505. 10.1016/J.MOLCEL.2005.03.032

Fujii, S., & Fuchs, R. P. (2004). Defining the position of the switches between replicative and bypass DNA polymerases. EMBO Journal, 23(21), 4342–4352. 10.1038/SJ.EMBOJ.7600438/ASSET/10D2F144-3829-430E-A30B-F2A2EF151E3E/ASSETS/GRAPHIC/EMBJ7600438-FIG-0005-M.JPG

Gallina, I., Hendriks, I. A., Hoffmann, S., Larsen, N. B., Johansen, J., Colding-Christensen, C. S., Schubert, L., Sellés-Baiget, S., Fábián, Z., Kühbacher, U., Gao, A. O., Räschle, M., Rasmussen, S., Nielsen, M. L., Mailand, N., & Duxin, J. P. (2021). The ubiquitin ligase RFWD3 is required for translesion DNA synthesis. Molecular Cell, 81(3), 442. 10.1016/J.MOLCEL.2020.11.029

Gersch, M., Gladkova, C., Schubert, A. F., Michel, M. A., Maslen, S., & Komander, D. (2017). Mechanism and regulation of the Lys6-selective deubiquitinase USP30. Nature Structural & Molecular Biology, 24(11), 920. 10.1038/NSMB.3475

Hershko, A., & Ciechanover, A. (1998). THE UBIQUITIN SYSTEM. Annual Review of Biochemistry, 67(1), 425–479. 10.1146/annurev.biochem.67.1.425

Hibbert, R. G., Huang, A., Boelens, R., & Sixma, T. K. (2011). E3 ligase Rad18 promotes monoubiquitination rather than ubiquitin chain formation by E2 enzyme Rad6. Proceedings of the National Academy of Sciences of the United States of America, 108(14), 5590–5595. 10.1073/pnas.1017516108

Hibbert, R. G., & Sixma, T. K. (2012a). Intrinsic Flexibility of Ubiquitin on Proliferating Cell Nuclear Antigen (PCNA) in Translesion Synthesis. Journal of Biological Chemistry, 287(46), 39216–39223. 10.1074/jbc.M112.389890

Hibbert, R. G., & Sixma, T. K. (2012b). Intrinsic Flexibility of Ubiquitin on Proliferating Cell Nuclear Antigen (PCNA) in Translesion Synthesis. Journal of Biological Chemistry, 287(46), 39216–39223. 10.1074/jbc.M112.389890

Hoege, C., Pfander, B., Moldovan, G. L., Pyrowolakis, G., & Jentsch, S. (2002a). RAD6-dependent DNA repair is linked to modification of PCNA by ubiquitin and SUMO. Nature 2002 419:6903, 419(6903), 135–141. 10.1038/nature00991

Hoege, C., Pfander, B., Moldovan, G. L., Pyrowolakis, G., & Jentsch, S. (2002b). RAD6-dependent DNA repair is linked to modification of PCNA by ubiquitin and SUMO. Nature 2002 419:6903, 419(6903), 135–141. 10.1038/nature00991

Hornbeck, P. V., Zhang, B., Murray, B., Kornhauser, J. M., Latham, V., & Skrzypek, E. (2015). PhosphoSitePlus, 2014: mutations, PTMs and recalibrations. Nucleic Acids Research, 43(Database issue), D512. 10.1093/NAR/GKU1267

Huang, T. T., Nijman, S. M. B., Mirchandani, K. D., Galardy, P. J., Cohn, M. A., Haas, W., Gygi, S. P., Ploegh, H. L., Bernards, R., & D’Andrea, A. D. (2006a). Regulation of monoubiquitinated PCNA by DUB autocleavage. Nature Cell Biology, 8(4), 341–347. 10.1038/ncb1378

Huang, T. T., Nijman, S. M. B., Mirchandani, K. D., Galardy, P. J., Cohn, M. A., Haas, W., Gygi, S. P., Ploegh, H. L., Bernards, R., & D’Andrea, A. D. (2006b). Regulation of monoubiquitinated PCNA by DUB autocleavage. Nature Cell Biology, 8(4), 341–347. 10.1038/ncb1378

Joosten, R. P., Long, F., Murshudov, G. N., & Perrakis, A. (2014a). The PDB_REDO server for macromolecular structure model optimization. IUCrJ, 1(Pt 4), 213–220. 10.1107/S2052252514009324

Joosten, R. P., Long, F., Murshudov, G. N., & Perrakis, A. (2014b). The PDB_REDO server for macromolecular structure model optimization. IUCrJ, 1(Pt 4), 213–220. 10.1107/S2052252514009324

Jumper, J., Evans, R., Pritzel, A., Green, T., Figurnov, M., Ronneberger, O., Tunyasuvunakool, K., Bates, R., Žídek, A., Potapenko, A., Bridgland, A., Meyer, C., Kohl, S. A. A., Ballard, A. J., Cowie, A., Romera-Paredes, B., Nikolov, S., Jain, R., Adler, J., … Hassabis, D. (2021). Highly accurate protein structure prediction with AlphaFold. Nature 2021 596:7873, 596(7873), 583–589. 10.1038/s41586-021-03819-2

Kannouche, P. L., Wing, J., & Lehmann, A. R. (2004). Interaction of Human DNA Polymerase η with Monoubiquitinated PCNA: A Possible Mechanism for the Polymerase Switch in Response to DNA Damage. Molecular Cell, 14(4), 491–500. 10.1016/S1097-2765(04)00259-X

Keijzer, N., Priyanka, A., Stijf-Bultsma, Y., Fish, A., Gersch, M., & Sixma, T. K. (2024). Variety in the USP deubiquitinase catalytic mechanism. Life Science Alliance, 7(4). 10.26508/LSA.202302533

Kirouac, K. N., & Ling, H. (2011). Unique active site promotes error-free replication opposite an 8-oxo-guanine lesion by human DNA polymerase iota. Proceedings of the National Academy of Sciences of the United States of America, 108(8), 3210–3215. 10.1073/PNAS.1013909108/SUPPL_FILE/PNAS.1013909108_SI.PDF

Komander, D., Clague, M. J., & Urbé, S. (2009). Breaking the chains: structure and function of the deubiquitinases. Nature Reviews Molecular Cell Biology, 10(8), 550–563. 10.1038/nrm2731

Komander, D., & Rape, M. (2012). The Ubiquitin Code. Annual Review of Biochemistry, 81(1), 203–229. 10.1146/annurev-biochem-060310-170328

Krijger, P. H. L., Lee, K.-Y., Wit, N., van den Berk, P. C. M., Wu, X., Roest, H. P., Maas, A., Ding, H., Hoeijmakers, J. H. J., Myung, K., & Jacobs, H. (2011). HLTF and SHPRH are not essential for PCNA polyubiquitination, survival and somatic hypermutation: existence of an alternative E3 ligase. DNA Repair, 10(4), 438–444. 10.1016/j.dnarep.2010.12.008

Lehmann, A. R., Niimi, A., Ogi, T., Brown, S., Sabbioneda, S., Wing, J. F., Kannouche, P. L., & Green, C. M. (2007). Translesion synthesis: Y-family polymerases and the polymerase switch. 10.1016/j.dnarep.2007.02.003

Li, H., Lim, K. S., Kim, H., Hinds, T. R., Jo, U., Mao, H., Weller, C. E., Sun, J., Chatterjee, C., D’Andrea, A. D., & Zheng, N. (2016). Allosteric Activation of Ubiquitin-Specific Proteases by β-Propeller Proteins UAF1 and WDR20. Molecular Cell, 63(2), 249–260. 10.1016/j.molcel.2016.05.031

Lim, K. S., Li, H., Roberts, E. A., Gaudiano, E. F., Clairmont, C., Sambel, L. A., Ponnienselvan, K., Liu, J. C., Yang, C., Kozono, D., Parmar, K., Yusufzai, T., Zheng, N., & D’Andrea, A. D. (2018). USP1 Is Required for Replication Fork Protection in BRCA1-Deficient Tumors. Molecular Cell, 72(6), 925–941.e4. 10.1016/j.molcel.2018.10.045

Mattiroli, F., Vissers, J. H. A., Van Dijk, W. J., Ikpa, P., Citterio, E., Vermeulen, W., Marteijn, J. A., & Sixma, T. K. (2012). RNF168 ubiquitinates K13-15 on H2A/H2AX to drive DNA damage signaling. Cell, 150(6), 1182–1195. 10.1016/j.cell.2012.08.005

Meng, E. C., Pettersen, E. F., Couch, G. S., Huang, C. C., & Ferrin, T. E. (2006). Tools for integrated sequence-structure analysis with UCSF Chimera. BMC Bioinformatics, 7(1), 1–10. 10.1186/1471-2105-7-339/TABLES/2

Mulder, M. P. C., El Oualid, F., Ter Beek, J., & Ovaa, H. (2014). A Native Chemical Ligation Handle that Enables the Synthesis of Advanced Activity-Based Probes: Diubiquitin as a Case Study. Chembiochem, 15(7), 946. 10.1002/CBIC.201402012

Nijman, S. M. B., Huang, T. T., Dirac, A. M. G., Brummelkamp, T. R., Kerkhoven, R. M., D’Andrea, A. D., & Bernards, R. (2005). The Deubiquitinating Enzyme USP1 Regulates the Fanconi Anemia Pathway. Molecular Cell, 17(3), 331–339. 10.1016/j.molcel.2005.01.008

Nijman, S. M. B., Luna-Vargas, M. P. A., Velds, A., Brummelkamp, T. R., Dirac, A. M. G., Sixma, T. K., & Bernards, R. (2005). A genomic and functional inventory of deubiquitinating enzymes. Cell, 123(5), 773–786. 10.1016/j.cell.2005.11.007

Nusawardhana, A., Pale, L. M., Nicolae, C. M., & Moldovan, G. L. (2024). USP1-dependent nucleolytic expansion of PRIMPOL-generated nascent DNA strand discontinuities during replication stress. Nucleic Acids Research, 52(5), 2340–2354. 10.1093/NAR/GKAD1237

Olazabal-Herrero, A., García-Santisteban, I., & Rodríguez, J. A. ntonio. (2016). Mutations in the ‘Fingers’ subdomain of the deubiquitinase USP1 modulate its function and activity. The FEBS Journal, 283(5), 929–946. 10.1111/FEBS.13648

Parker, J. L., & Ulrich, H. D. (2009). Mechanistic analysis of PCNA poly-ubiquitylation by the ubiquitin protein ligases Rad18 and Rad5. The EMBO Journal, 28(23), 3657–3666. 10.1038/emboj.2009.303

Pettersen, E. F., Goddard, T. D., Huang, C. C., Meng, E. C., Couch, G. S., Croll, T. I., Morris, J. H., & Ferrin, T. E. (2021). UCSF ChimeraX: Structure visualization for researchers, educators, and developers. Protein Science : A Publication of the Protein Society, 30(1), 70. 10.1002/PRO.3943

Piberger, A. L., Bowry, A., Kelly, R. D. W., Walker, A. K., González-Acosta, D., Bailey, L. J., Doherty, A. J., Méndez, J., Morris, J. R., Bryant, H. E., & Petermann, E. (2020). PrimPol-dependent single-stranded gap formation mediates homologous recombination at bulky DNA adducts. Nature Communications 2020 11:1, 11(1), 1–14. 10.1038/s41467-020-19570-7

Pichler, A., Knipscheer, P., Oberhofer, E., Van Dijk, W. J., Körner, R., Olsen, J. V., Jentsch, S., Melchior, F., & Sixma, T. K. (2005). SUMO modification of the ubiquitin-conjugating enzyme E2-25K. Nature Structural & Molecular Biology 2005 12:3, 12(3), 264–269. 10.1038/nsmb903

Pickart, C. M., & Raasi, S. (2005). Controlled Synthesis of Polyubiquitin Chains. Methods in Enzymology, 399, 21–36. 10.1016/S0076-6879(05)99002-2

Punjani, A., & Fleet, D. J. (2023). 3DFlex: determining structure and motion of flexible proteins from cryo-EM. Nature Methods 2023 20:6, 20(6), 860–870. 10.1038/s41592-023-01853-8

Punjani, A., Rubinstein, J. L., Fleet, D. J., & Brubaker, M. A. (2017). cryoSPARC: algorithms for rapid unsupervised cryo-EM structure determination. Nature Methods, 14(3), 290–296. 10.1038/NMETH.4169

Rennie, M. L., Arkinson, C., Chaugule, V. K., Toth, R., & Walden, H. (2021a). Structural basis of FANCD2 deubiquitination by USP1−UAF1. Nature Structural and Molecular Biology, 28(4), 356–364. 10.1038/s41594-021-00576-8

Rennie, M. L., Gundogdu, M., Arkinson, C., Liness, S., Frame, S., & Walden, H. (2024). Structural and Biochemical Insights into the Mechanism of Action of the Clinical USP1 Inhibitor, KSQ-4279. Journal of Medicinal Chemistry, 67(17), 15557–15568. 10.1021/ACS.JMEDCHEM.4C01184/SUPPL_FILE/JM4C01184_SI_001.CSV

Sale, J. E., Lehmann, A. R., & Woodgate, R. (2012). Y-family DNA polymerases and their role in tolerance of cellular DNA damage. Nature Reviews Molecular Cell Biology, 13(3), 141–152. 10.1038/nrm3289

Sanchez-Garcia, R., Gomez-Blanco, J., Cuervo, A., Carazo, J. M., Sorzano, C. O. S., & Vargas, J. (2021). DeepEMhancer: a deep learning solution for cryo-EM volume post-processing. Communications Biology 2021 4:1, 4(1), 1–8. 10.1038/s42003-021-02399-1

Sato, Y., Goto, E., Shibata, Y., Kubota, Y., Yamagata, A., Goto-Ito, S., Kubota, K., Inoue, J. I., Takekawa, M., Tokunaga, F., & Fukai, S. (2015). Structures of CYLD USP with Met1- or Lys63-linked diubiquitin reveal mechanisms for dual specificity. Nature Structural & Molecular Biology 2015 22:3, 22(3), 222–229. 10.1038/nsmb.2970

Simoneau, A., Engel, J. L., Bandi, M., Lazarides, K., Liu, S., Meier, S. R., Choi, A. H., Zhang, H., Shen, B., Martires, L., Gotur, D., Pham, T. V., Li, F., Gu, L., Gong, S., Zhang, M., Wilker, E., Pan, X., Whittington, D. A., … Feng, T. (2023). Ubiquitinated PCNA Drives USP1 Synthetic Lethality in Cancer. Molecular Cancer Therapeutics, 22(2), 215–226. 10.1158/1535-7163.MCT-22-0409/709732/AM/UBIQUITINATED-PCNA-DRIVES-USP1-SYNTHETIC-LETHALITY

Stelter, P., & Ulrich, H. D. (2003). Control of spontaneous and damage-induced mutagenesis by SUMO and ubiquitin conjugation. Nature, 425(6954), 188–191. 10.1038/NATURE01965

Tirman, S., Quinet, A., Wood, M., Meroni, A., Cybulla, E., Jackson, J., Pegoraro, S., Simoneau, A., Zou, L., & Vindigni, A. (2021). Temporally distinct post-replicative repair mechanisms fill PRIMPOL-dependent ssDNA gaps in human cells. Molecular Cell, 81(19), 4026–4040.e8. 10.1016/J.MOLCEL.2021.09.013/ATTACHMENT/359B963E-864A-4443-A3E5-54C1CF508A8D/MMC1.PDF

Tsaalbi-Shtylik, A., Mingard, C., Räz, M., Oka, R., Manders, F., Van Boxtel, R., De Wind, N., & Sturla, S. J. (2024). DNA mismatch repair controls the mutagenicity of Polymerase ζ-dependent translesion synthesis at methylated guanines. DNA Repair, 142, 103755. 10.1016/J.DNAREP.2024.103755

Unk, I., Hajdú, I., Fátyol, K., Hurwitz, J., Yoon, J.-H., Prakash, L., Prakash, S., & Haracska, L. (2008). Human HLTF functions as a ubiquitin ligase for proliferating cell nuclear antigen polyubiquitination. Proceedings of the National Academy of Sciences of the United States of America, 105(10), 3768– 3773. 10.1073/pnas.0800563105

Unk, I., Hajdú, I., Fátyol, K., Szakál, B., Blastyák, A., Bermudez, V., Hurwitz, J., Prakash, L., Prakash, S., & Haracska, L. (2006). Human SHPRH is a ubiquitin ligase for Mms2-Ubc13-dependent polyubiquitylation of proliferating cell nuclear antigen. Proceedings of the National Academy of Sciences of the United States of America, 103(48), 18107–18112. 10.1073/PNAS.0608595103

Yang, K., Moldovan, G.-L., Vinciguerra, P., Murai, J., Takeda, S., & D’Andrea, A. D. (2011). Regulation of the Fanconi anemia pathway by a SUMO-like delivery network. Genes & Development, 25(17), 1847–1858. 10.1101/gad.17020911

Yang, X. H., & Zou, L. (2009). Dual functions of DNA replication forks in checkpoint signaling and PCNA ubiquitination. Cell Cycle, 8(2), 191–194. 10.4161/cc.8.2.7357

Ye, Y., Akutsu, M., Reyes-Turcu, F., Enchev, R. I., Wilkinson, K. D., & Komander, D. (2011). Polyubiquitin binding and cross-reactivity in the USP domain deubiquitinase USP21. EMBO Reports, 12(4), 350–357. 10.1038/EMBOR.2011.17/SUPPL_FILE/EMBR201117-SUP-0001.PDF

Yin, J., Schoeffler, A. J., Wickliffe, K., Newton, K., Starovasnik, M. A., Dueber, E. C., & Harris, S. F. (2015). Structural Insights into WD-Repeat 48 Activation of Ubiquitin-Specific Protease 46. Structure, 23(11), 2043–2054. 10.1016/j.str.2015.08.010

Zhu, H., Zhang, T., Wang, F., Yang, J., & Ding, J. (2019). Structural insights into the activation of USP46 by WDR48 and WDR20. Cell Discovery 2019 5:1, 5(1), 1–4. 10.1038/s41421-019-0102-1

