## supplemental figures and tables for "USP1/UAF1 targets polyubiquitinated PCNA with an exo-cleavage mechanism that enriches for monoubiquitinated PCNA"

**Supplementary Figure 1 –Ubiquitinated PCNA substrates used for experiments**

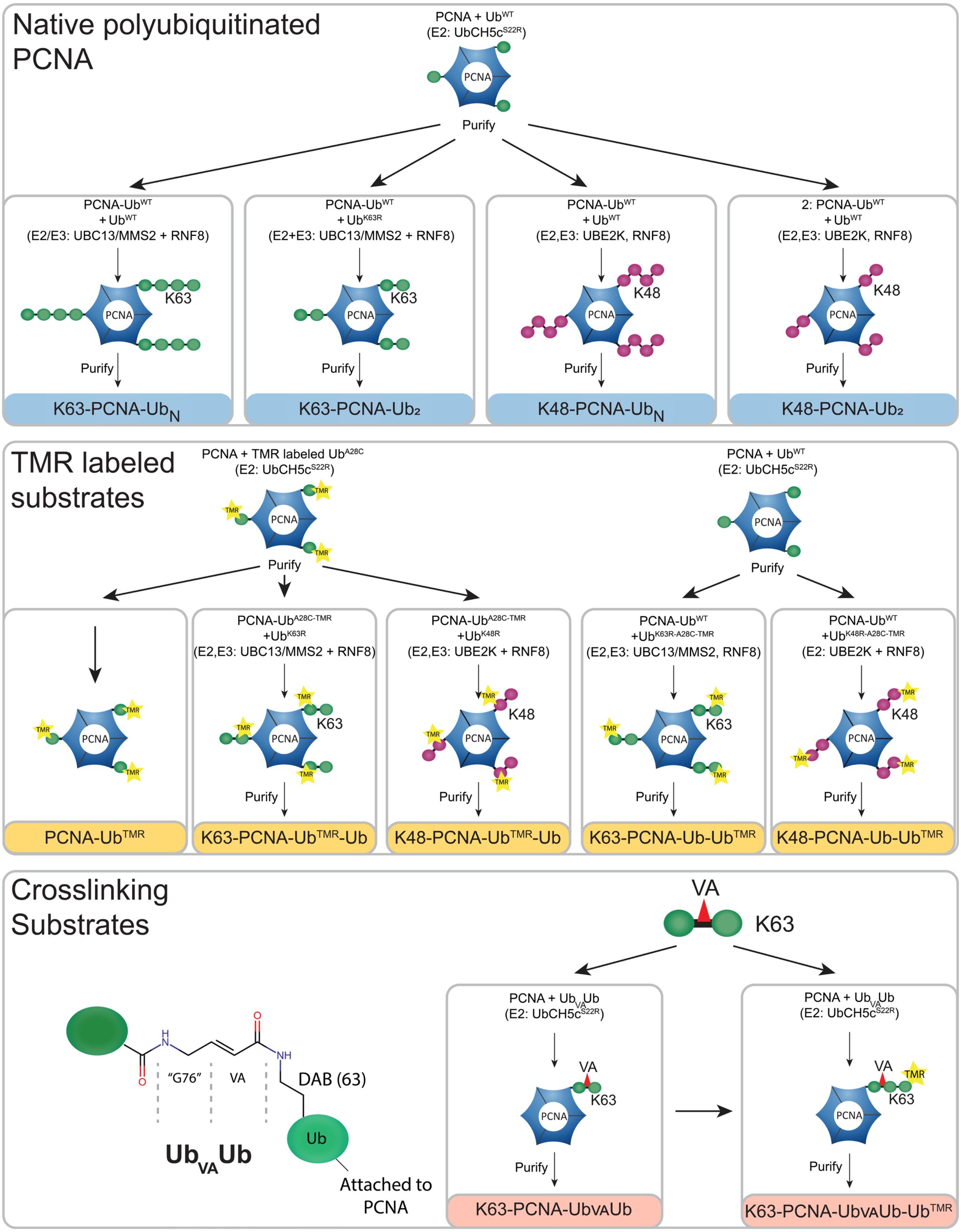

**Supplementary figure 2 – A TAMRA label on residue 28 (A28C) of ubiquitin does not interfere with reactions**
**A)** DUB assay of USP1/UAF1 on PCNA-Ub and PCNA-Ub$^{TMR}$. Coomassie staining (left panel) shows that TMR-labeled Ub$^{A28C}$ does not hinder deubiquitination by USP1/UAF1. **B)** Fluorescent scan of the PCNA-Ub$^{TMR}$ DUB assay. **C)** DUB assay on K63-PCNA-Ub-Ub compared with K63-PCNA-Ub$^{TMR}$-Ub and K63-PCNA-Ub-Ub$^{TMR}$ shows that the Ub$^{A28C}$ label does not affect cleavage. **D)** Fluorescence image of gel shown in C. **E)** DUB assays on K48-PCNA-Ub-Ub compared with A28C labeled K48-PCNA-Ub$^{TMR}$-Ub and K48-PCNA-Ub-Ub$^{TMR}$ shows that the label does not affect cleavage. **F)** Fluorescence image of gel shown in F.

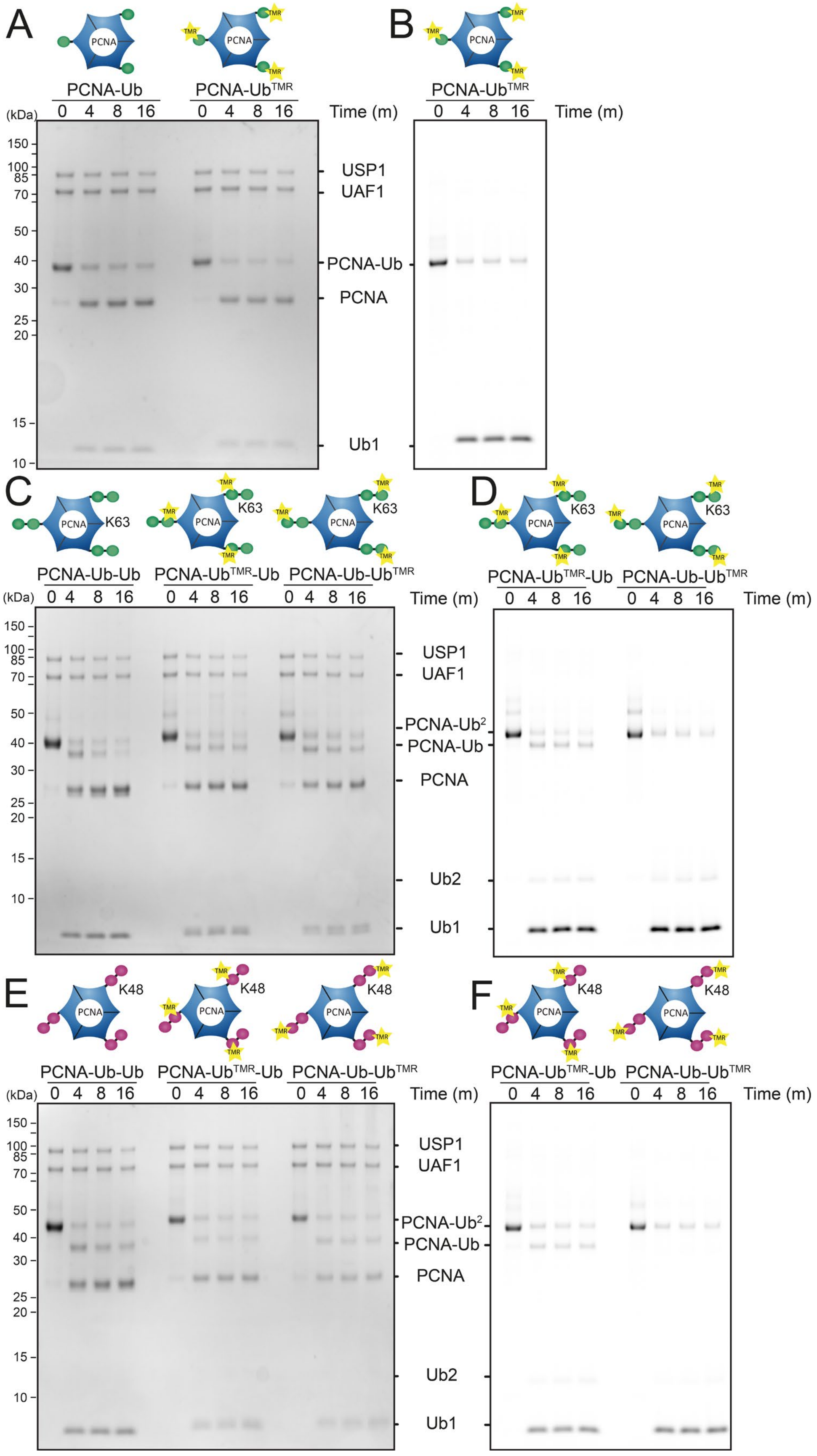

**Supplementary figure 3 – Experimental data used for kinetic modeling**

**A)** FP assays on PCNA-Ub$^{TMR}$, proximally and distally labeled K63-PCNA-Ub$_2$ A single concentration of USP1 (10 nM) was tested against four concentrations (4 µM, 2 µM, 1 µM, 0.5 µM) of labeled substrate. **B)** FP assays on proximally and distally labeled K48-PCNA-Ub$_2$. **C)** MST analysis of USP1$^{WT}$/UAF1$^{WT}$ binding to PCNA-Ub$^{L73P}$, PCNA and ubiquitin. For this analysis, USP1/UAF1 was labeled using DY-574P1-maleimide (methods). **D)** Ub$^{Rho}$ fluorescence intensity assay performed in previous publication (Keijzer et al., 2024; Fig. S2A, upper left panel) was used to accurately determine $K_{cut}$ (cleavage) to support global kinetic analysis.

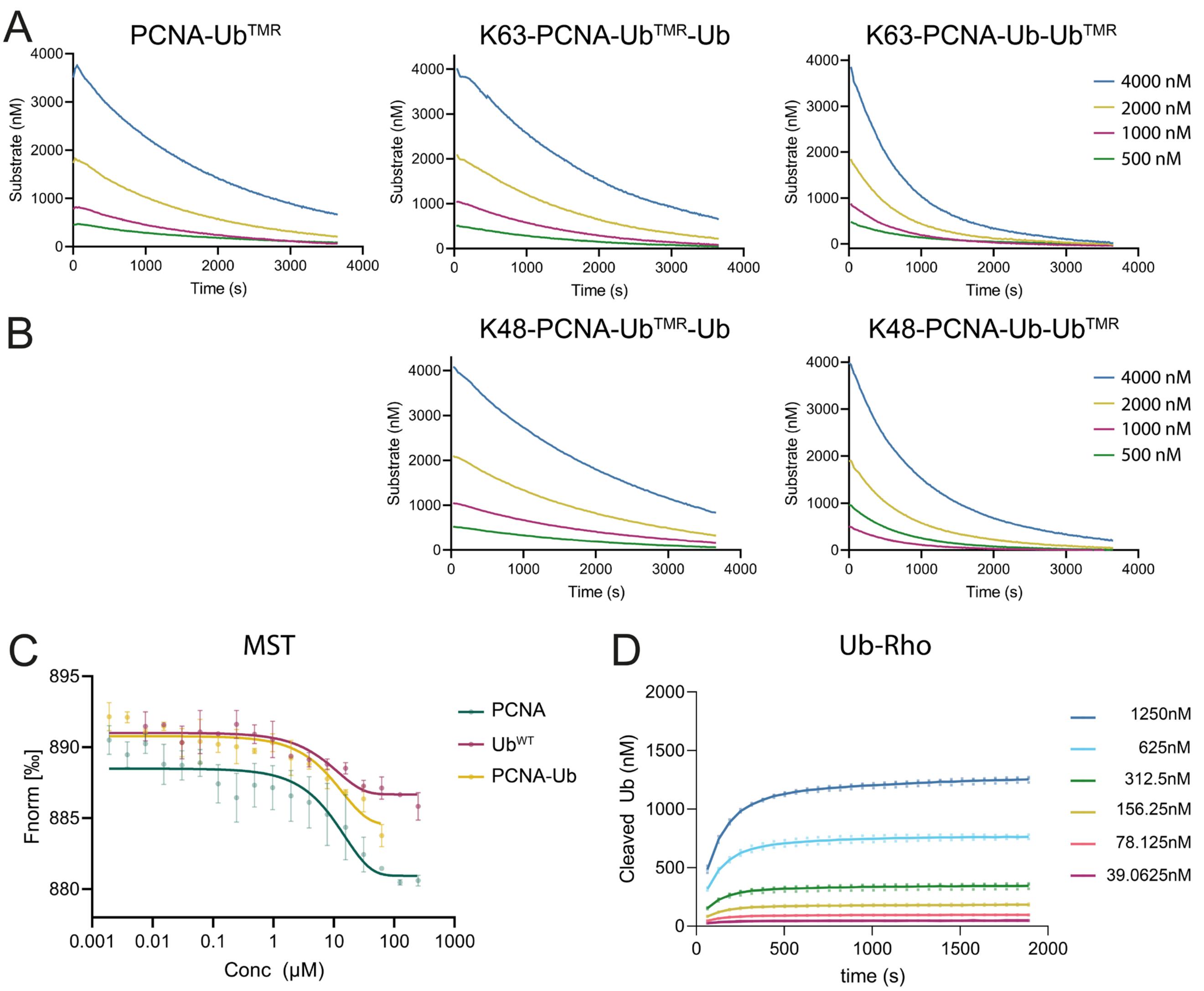

**Supplementary figure 4 – Output of kinetic model** **A)** Kinetic fits generated using Kintek model for K63 substrates (grey lines) plotted against FP data shown in supplementary figure 2 (dots). For PCNA-Ub$^{TMR}$ and K63-PCNA-Ub-Ub$^{TMR}$ the 500 nM data was excluded from the kinetic model. For K48-PCNA-Ub-Ub$^{TMR}$ the 2000nM data was excluded from the kinetic model **B)** Kinetic fits generated using Kintek model for K48 substrates (grey lines) plotted against experimental data (dots). For both substrates, 2000 nM data was excluded from the kinetic model. **C)** Michaelis-Menten analysis of distally labeled K63- and K48-PCNA-$Ub_2$ and PCNA-$Ub_2$ using data from the kinetic model. **D)** Substrate inhibition analysis of proximally labeled K63- and K48-PCNA-$Ub_2$ used to calculate reaction rates, using data from the kinetic model. **E)** Simulation by kinetic model for K63-PCNA-Ub$^{TMR}$-Ub (same as in Fig. 2C) and K63-PCNA-Ub-Ub$^{TMR}$ shows cleavage of substrates and appearance of different products as USP1/UAF1 cleaves PCNA-$Ub_2$. **F)** Same as in E) but for K48-PCNA-Ub$^{TMR}$-Ub and K48-PCNA-Ub-Ub$^{TMR}$.

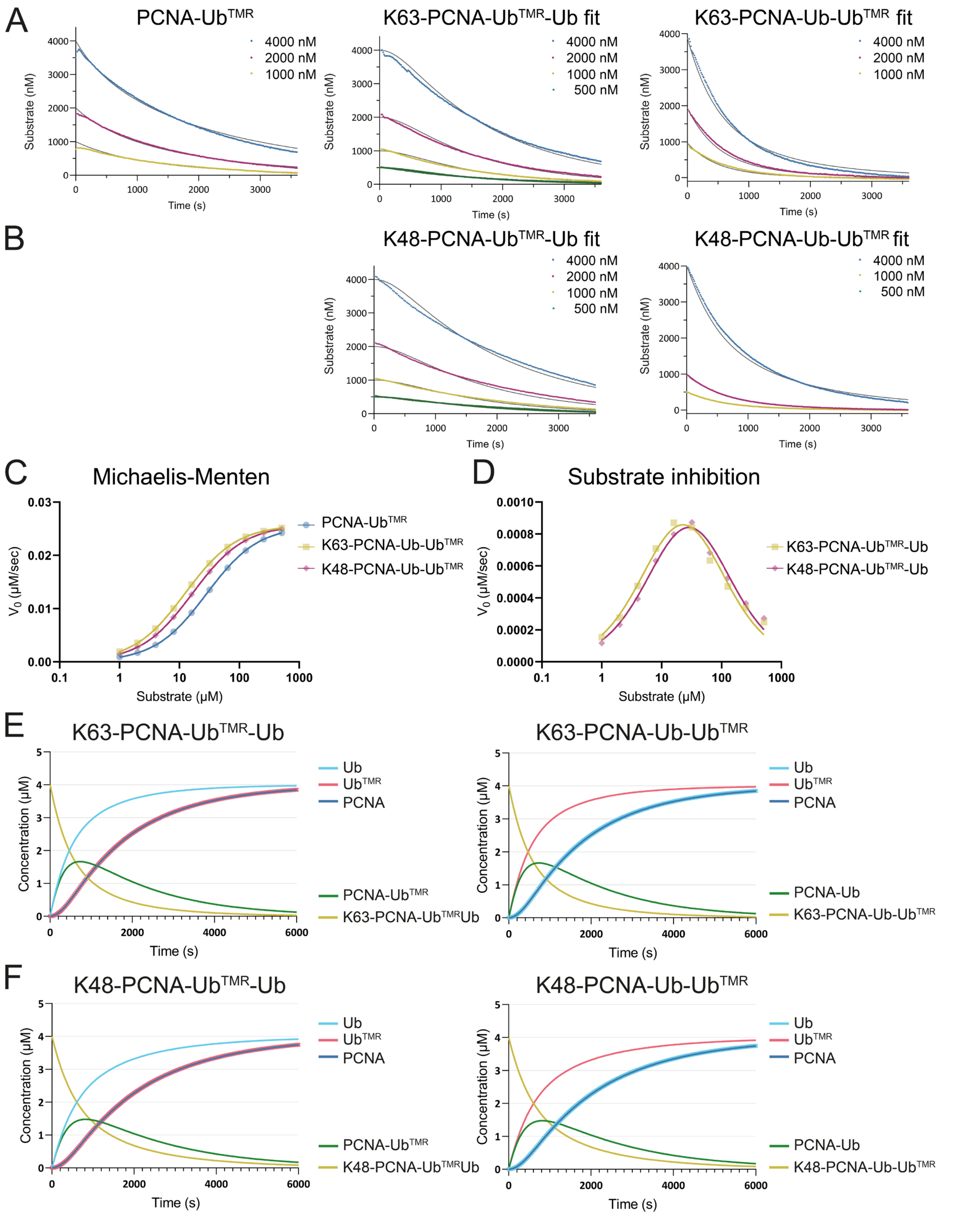

### Supplementary figure 5 – Cryo-EM data processing workflow

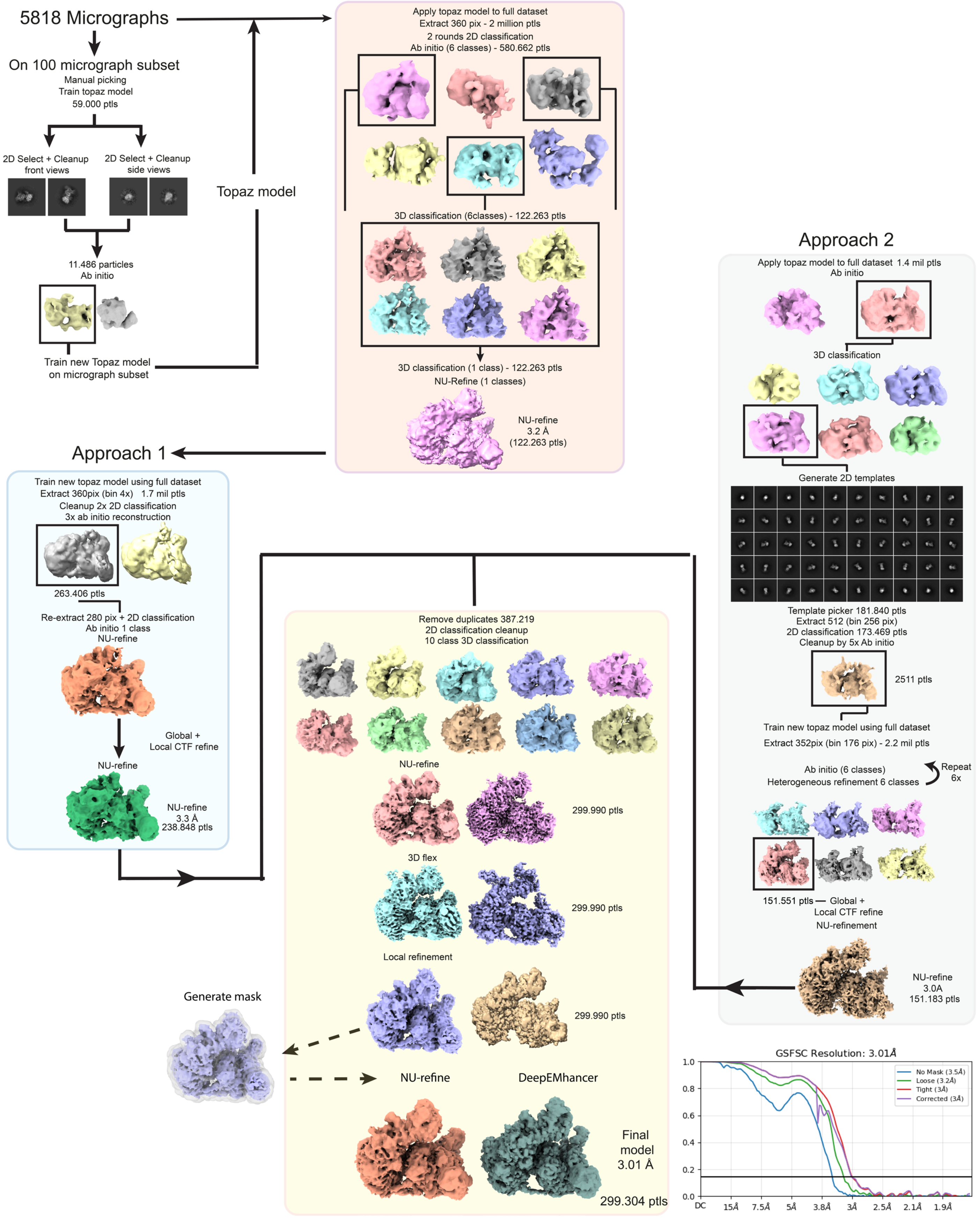

**Supplementary figure 6 – Positioning of the ABU-linker in USP1s active site**

**A)** Structure of USP1/UAF1 covalently bound to K63-Ub$_{VA}$Ub. **B)** Zoom on the USP1 active site, showing catalytic C90 covalently bound to the formed 4-aminobutyric acid (ABU) linker (dark green); the ABU linker now connects G75 of the distal ubiquitin (dark green) with a 2,4-diaminobutyric acid residue (DAB) at position 63 of the proximal ubiquitin (light green) (see D). Catalytic residues C90, H593 and D751 and D752 are highlighted. **C)** Insert3 of USP1 anchors on the WD40 domain of UAF1 and reveals a new binding site facilitated by three different Hydrogen bonds. **D)** Schematic of the K63-Ub$_{VA}$Ub linkage before and after conjugation to USP1's C90. As shown in the bottom panel, the K63-Ub$_{VA}$Ub linkage mimics the native isopeptide linkage between G76 of the distal ubiquitin and K63 of the proximal ubiquitin**. E)** D752 is essential for catalysis in USP1 (Keijzer et al,. 2024), but the density (lower panel) surrounding this residue allows for two different rotamers. We followed the canonical positioning (pink) in the deposited coordinates.

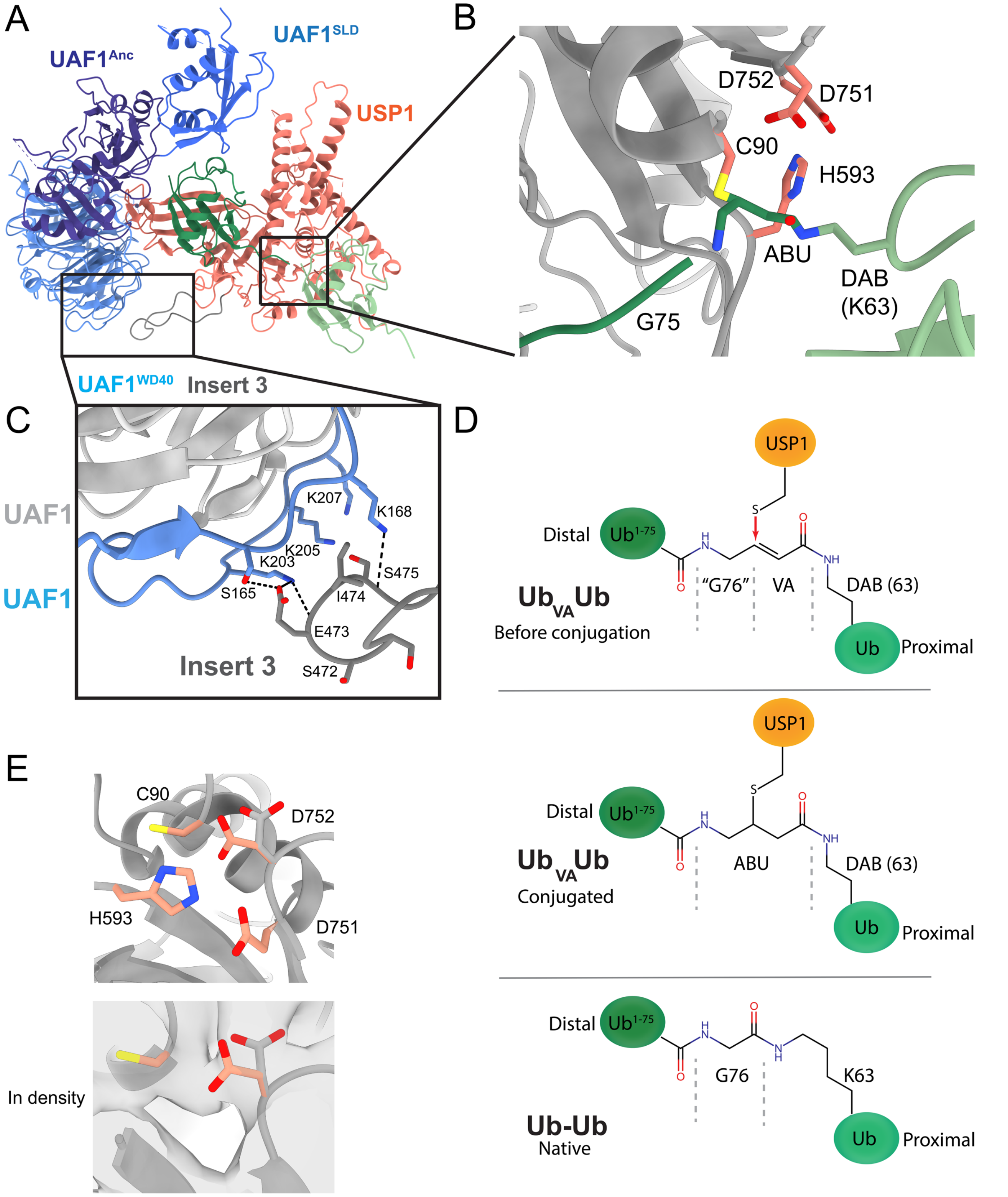

**Supplementary figure 7 – Stability of UAF1 deletion mutants is comparible to UAF1$^{WT}$**

Thermal stability analysis of UAF1 variants show only slight difference in melting temperature compared with UAF1$^{WT}$. Inflection points are shown in Supplementary table 3.

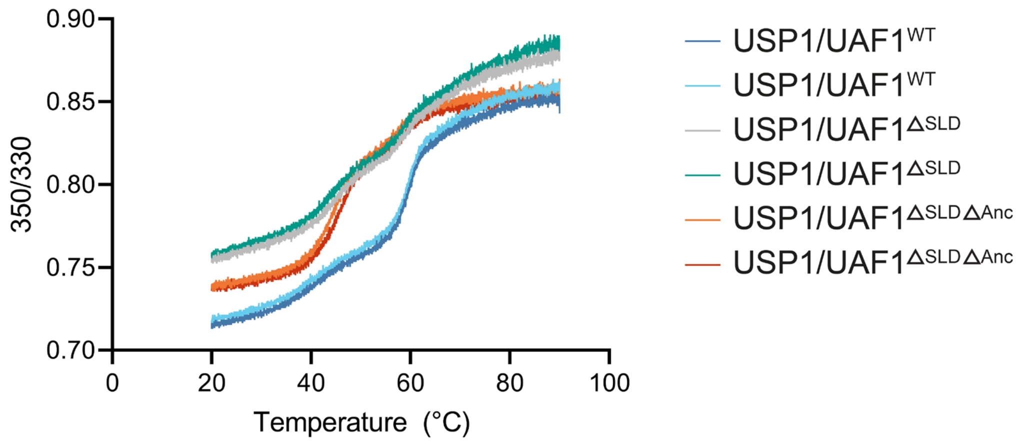

**Supplementary figure 8 – Hyperactivation of USP1/UAF1 by mutating a conserved negative patch.**

**A)** Superposition of D172/E173 loop of USP1/UAF1, compared with equivalent loop in USP30 and their AlphaFold2 predictions shows that the loop is correctly predicted. Other human USPs do not have this loop, and are therefore not shown. Structure solved in this paper is solved in pink, superposed on USP1 AlphaFold2 prediction (grey). USP30 (5OHP (Gersch et al., 2017)) shown in blue with its AlphaFold2 prediction in grey. **B)** Superposition of USP1 and USP30 (5OHP) showing interface with the proximal ubiquitin. The D172/E173 equivalent residues in USP30 (D134/D135) are positioned further from the proximal ubiquitin. More importantly, since USP30 binds a K6-linked diubiquitin chain and as such the proximal ubiquitin is positioned different from how USP1 interacts the K63 proximal ubiquitin. **C)** PAE plots of USP1 and USP30 AlphaFold2 predictions.

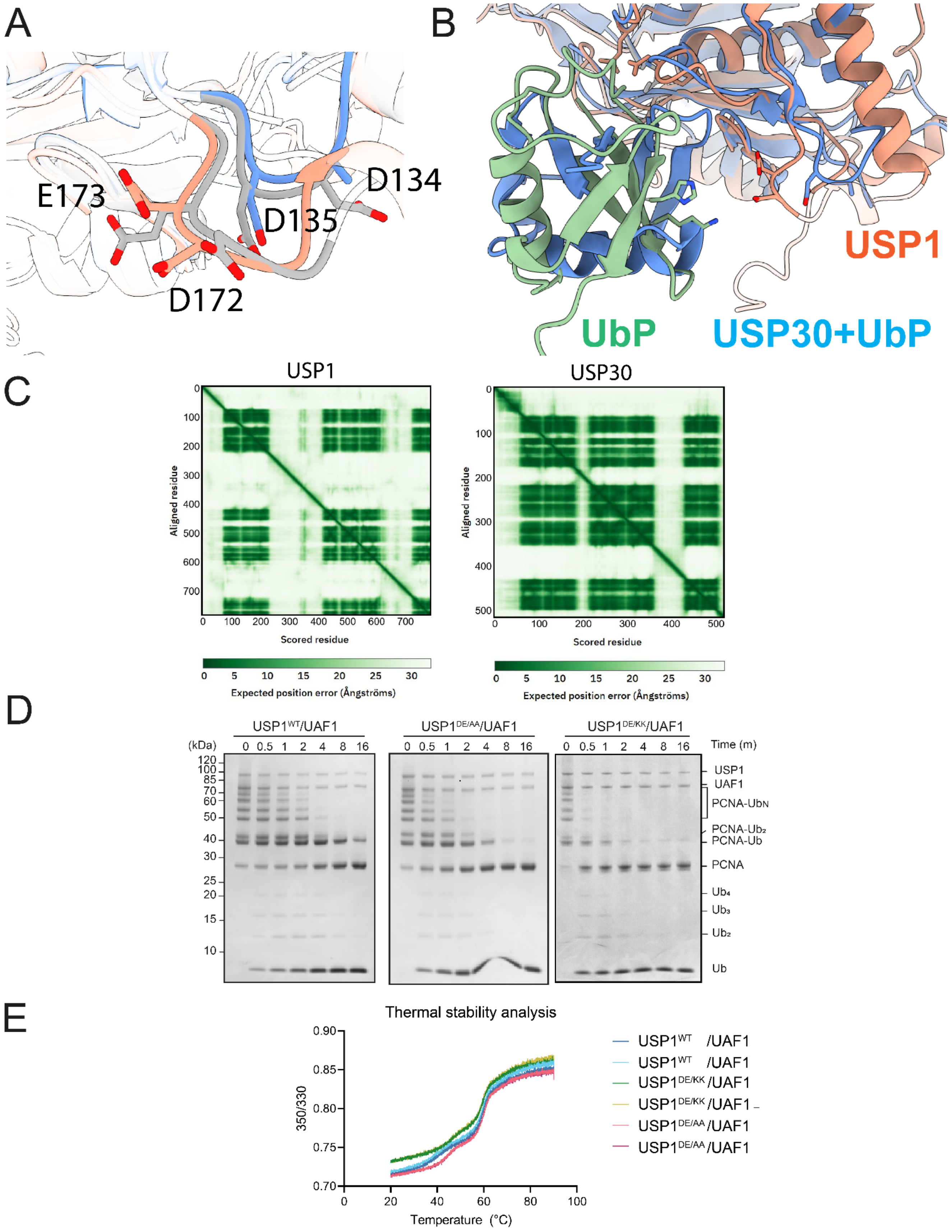

**D)** DUB assay on K63-PCNA-$Ub_N$ comparing $USP1^{WT}$/UAF1, $USP1^{DE/AA}$/UAF1 $USP1^{DE/KK}$/UAF1. Mutating D172 and E173 causes hyper-activation of USP1 and does not change its cleavage mechanism. **E)** Thermal stability assay on $USP1^{D172/E173}$/UAF1 variants. $USP1^{DE/KK}$ and $USP1^{DE/AA}$ have higher thermal stability compared with $USP1^{WT}$. Inflection points are shown in Table S3.

**Supplementary table 1 - Kinetic model and parameters/constants**

Constants describing the different reactions and equilibriums in the kinetic models. $K_{on}$ is diffusion limited and therefore the same for each reaction. $k_1$, $k_2$, $k_3$, $k_4$, $k_5$, $k_6$ are the off rates for each reaction and can be seen as representations of affinity in this model. Initially, the kinetic constants for each K63- or K48-specific step were defined as independent, except the cleavage constant ($k_{on}$ *and* $k_{cut}$). However, after fitting we realized that all constants for both linkages, except for $k_1$, are similar, and thus were shared in the subsequent fitting steps. Additionally, dissociation rates of PCNA_USP1_Ub and PCNA_USP1 were originally independent from each other, but appeared to be similar and were defined as a single constant ($k_6$) to describe both.

$$USP1 + PCNA\_Ub\_Ub \underset{k_1}{\overset{k_{on}}{\rightleftharpoons}} PCNA\_Ub\_Ub\_USP1 \overset{k_{cut}}{\rightarrow} PCNA\_Ub\_USP1\_Ub \underset{k_{on}}{\overset{k_2}{\rightleftharpoons}} PCNA\_Ub + USP1\_Ub \underset{k_{on}}{\overset{k_3}{\rightleftharpoons}} USP1 + Ub$$

$$PCNA\_Ub\_Ub\_USP1 \overset{k_4}{\rightarrow} PCNA\_Ub\_USP1 + Ub$$

$$USP1 + PCNA\_Ub \underset{k_5}{\overset{k_{on}}{\rightleftharpoons}} PCNA\_Ub\_USP1 \overset{k_{cut}}{\rightarrow} PCNA\_USP1\_Ub \underset{k_{on}}{\overset{k_6}{\rightleftharpoons}} PCNA + USP1\_Ub \underset{k_{on}}{\overset{k_3}{\rightleftharpoons}} USP1 + Ub$$

$$USP1 + PCNA \underset{k_6}{\overset{k_{on}}{\rightleftharpoons}} PCNA\_USP1$$

| | **K63 kinetic model** | **K48 kinetic model** |
|---|---|---|
| $k_{on}$ **(µM$^{-1}$*s$^{-1}$)** | 100 | 100 |
| $k_{cut}$ **(s$^{-1}$)** | 2.6 ± 0.06 | 2.6 ± 0.06 |
| $k_1$ **(s$^{-1}$)** | 1244 ± 32 | 1656 ± 32 |
| $k_2$ **(s$^{-1}$)** | 3840 ± 485 | 3840 ± 485 |
| $k_3$ **(s$^{-1}$)** | 932 ± 554 | 932 ± 554 |
| $k_4$ **(s$^{-1}$)** | 5.5 ± 3 | 5.5 ± 3 |
| $k_5$ **(s$^{-1}$)** | 2812 ± 60 | 2812 ± 60 |
| $k_6$ **(s$^{-1}$)** | 229 ± 6.7 | 229 ± 6.7 |

**Supplementary table 2 – Cryo-EM data collection and refinement statistics**

| **Data collection and processing** | **EMD:** 52316 |
|---|---|
| Magnification | 105.000x |
| Voltage (kV) | 300 |
| Electron exposure (e–/Å$^2$) | 50 |
| Defocus range (μm) | -1.2, -2.4 |
| Pixel size (Å) | 0.836 |
| Symmetry imposed | C1 |
| Initial particle images (no.) | 3.110.350 |
| Final particle images (no.) | 299.304 |
| Map resolution (Å)<br>FSC threshold | 3.01<br>0.143 |
| Map sharpening *B* factor (Å$^2$) | DeepEMhancer |
| **Refinement** | **PDB:** 9NHW |
| Initial model used<br>(PDB code) | Alphafold2<br>1UBQ |
| Model resolution (Å)<br>FSC threshold | 3.47<br>0.143 |
| Model composition<br>Non-hydrogen atoms<br>Protein residues<br>Ligands | <br>9350<br>1180<br>3 |
| *B* factors (Å$^2$)<br>Protein | <br>116 |
| R.m.s. deviations<br>Bond lengths (Å)<br>Bond angles (°) | <br>0.004 (3)<br>0.555 (2) |
| Validation<br>MolProbity score<br>Clashscore<br>Poor rotamers (%) | <br>1.73<br>9.43<br>0.19 |
| Ramachandran plot<br>Favored (%)<br>Allowed (%)<br>Disallowed (%) | <br>96.57<br>3.43<br>0.00 |

**Supplementary table 3 – Thermal stability of USP1/UAF1 mutants.**

Inflection points of USP1/UAF1 variants measured using microscale thermophoresis. USP1/UAF1 has two inflection points for USP1 (#1) and UAF1 (#2). Interestingly, double mutants of $USP1^{D172/E173}$ causes an increase in thermal stability. Deletions of UAF1 only cause minor decrease in thermal stability.

| | **Inflection point #1 (USP1)** | **Inflection point #2 (UAF1)** |
|---|---|---|
| $USP1^{wt}$ /UAF1 | 41.8 °C | 59.7 °C |
| $USP1^{wt}$ /UAF1 | 42.9 °C | 59.3 °C |
| $USP1^{DE/KK}$/UAF1 | 46.3 °C | 59.5 °C |
| $USP1^{DE/KK}$/UAF1 | 46.4 °C | 59.5 °C |
| $USP1^{DE/AA}$/UAF1 | 44.03 °C | 60.1 °C |
| $USP1^{DE/AA}$/UAF1 | 45.1 °C | 60.2 °C |
| $USP1^{WT}$ /$UAF1^{\Delta SLD}$ | 44.8 °C | 59.3 °C |
| $USP1^{WT}$ /$UAF1^{\Delta SLD}$ | 44.2 °C | 58.5 °C |
| $USP1^{WT}$ /$UAF1^{\Delta ANC+SLD}$ | 44.7 °C | 57.1 °C |
| $USP1^{WT}$ /$UAF1^{\Delta ANC+SLD}$ | 45.8 °C | 58.5 °C |
